# NOTCH-mediated non-cell autonomous regulation of chromatin structure during senescence

**DOI:** 10.1101/212829

**Authors:** Aled J. Parry, Matthew Hoare, Dóra Bihary, Robert Hänsel-Hertsch, Stephen Smith, Kosuke Tomimatsu, Shankar Balasubramanian, Hiroshi Kimura, Shamith A. Samarajiwa, Masashi Narita

**Affiliations:** University of Cambridge, Cancer Research UK Cambridge Centre, Robinson Way, Cambridge, CB2 0RE, UK; University of Cambridge, Department of Medicine, Addenbrooke’s Hospital, Cambridge, CB2 0QQ, UK.; University of Cambridge, MRC Cancer Cell Unit, Hutchison / MRC Research Centre, Cambridge Biomedical; University of Cambridge, Department of Pathology, Addenbrooke’s Hospital, Cambridge, CB2 0QQ, UK. Campus, Cambridge, CB2 0XZ, UK; Department of Chemistry, University of Cambridge, Lensfield Road, Cambridge CB2 1EW, UK.; Tokyo Institute of Technology, Yokohama, 226-8501, Japan

## Abstract

Senescent cells interact with the surrounding microenvironment achieving diverse functional outcomes. In addition to autocrine and paracrine signalling mediated by factors of the senescence-associated secretory phenotype, we have recently identified that NOTCH1 can drive ‘lateral induction’ of a unique form of senescence in adjacent cells through specific induction of the NOTCH ligand JAG1. Here we show that NOTCH signalling can modulate chromatin structure both autonomously and non-autonomously. In addition to senescence-associated heterochromatic foci (SAHF), oncogenic RAS-induced senescent (RIS) cells in culture exhibit a massive increase in nucleosome-free regions (NRFs). NOTCH signalling suppresses both SAHF and NFR formation in this context. Strikingly, NOTCH-induced senescent cells, or cancer cells with high JAG1 expression, also drive similar chromatin architectural changes in adjacent cells through cell-cell contact. Mechanistically, we show that NOTCH signalling represses the chromatin architectural protein HMGA1, an association found in a range of human cancers. Thus, HMGA1 is involved not only in SAHFs, but also RIS-specific NFR formation. In conclusion, this study identifies that the JAG1-NOTCH-HMGA1 axis mediates the juxtacrine regulation of chromatin architecture.

## INTRODUCTION

Cellular senescence is an autonomous tumour suppressor mechanism that can be triggered by multiple pathophysiological stimuli including replicative exhaustion ^1,2^, exposure to chemotherapeutic drugs ^3^ and hyper-activation of oncogenes such as RAS ^4,5^. Persistent cell cycle arrest, an essential feature that defines senescence, is accompanied by diverse transcriptional, biochemical, and morphological alterations. These senescence hallmarks include increased expression and secretion of soluble factors (senescence-associated secretory phenotype; SASP) ^6,7^, senescence-associated β-galactosidase activity, enlarged nuclei, and dramatic alterations to chromatin structure ^8-10^. Importantly, the combination, quantity and quality of these features can vary depending on the type of senescence. Senescent cells have profound non-cell autonomous functionality mediated primarily by the SASP. The SASP can have either pro-or anti-tumorigenic effects and act in an autocrine or paracrine fashion ^11-14^. In addition, we have recently identified that NOTCH signalling can drive a cell-contact dependent juxtacrine senescence ^15^.

The NOTCH signalling pathway is involved in a wide array of developmental and physiological processes. During development and in tissue homeostasis NOTCH has well defined roles in differentiation and stem cell fate ^16^. Perturbations have been linked to tumorigenesis where, depending on the tissue of origin and context, NOTCH can have either oncogenic or tumour suppressive functionality ^17^. The pathway involves proteolytic cleavage of the NOTCH receptor upon contact mediated activation by a ligand of the JAGGED (JAG) or DELTA family on the surface of an adjacent cell. The cleaved NOTCH-intracellular domain (N-ICD) translocates to the nucleus where, together with transcriptional co-activators such as mastermind-like 1 (MAML1), it drives transcription of canonical target genes including the HES / HEY family of transcription factors ^16^. NOTCH signaling has also been shown to induce a type of senescence, NOTCH-induced senescence (NIS), where cells are characterised by distinct SASP components ^15,18^. Recently, we showed that during NIS there is a dramatic and specific upregulation of JAG1 that can activate NOTCH1 signaling and drive NIS in adjacent cells (called ‘lateral induction’) ^15^.

During senescence, particularly in oncogenic RAS-induced senescent (RIS) fibroblasts, there are characteristic changes to chromatin that culminate in the formation of senescence-associated heterochromatic foci (SAHFs) ^19,20^; layered structures facilitated by spatial rearrangement of existing heterochromatin ^21^. Other alterations include a loss of Lamin B1 from the nuclear periphery ^22-25^ and a decondensation of satellite heterochromatin (senescence-associated distention of satellites, SADS) ^26^.

SAHF formation is dependent on the transcriptional upregulation and accumulation of chromatin-bound High-Mobility Group A (HMGA) proteins ^27^. These are a family of flexible architectural proteins, consisting of HMGA1 and HMGA2, which bind to the minor groove of AT-rich DNA via three AT-hook domains to alter chromatin structure ^28,29^. Despite a critical role in the formation of SAHFs during senescence, HMGA proteins are also important during development where they promote tissue growth ^30,31^ and regulate differentiation ^32-35^. Furthermore, a large body of literature has accumulated that in most cases demonstrate an association between high HMGA expression and tumour malignancy as indicated by metastasis, chemo-resistance and poor prognosis ^36,37^. For example, HMGAs are upregulated in approximately 90% of lung carcinomas ^38^.

Chromatin accessibility at regulatory elements including promoters and enhancers is critical in determining gene expression profiles and is highly correlated with biological activity ^39,40^. Differentiated cells have a distribution of accessible regions characteristic of lineage and function, where more similar cell types share more open chromatin sites ^41^. The proportion of sites shared with embryonic stem cells (ESCs) is indicative of developmental maturity, as this is reduced as differentiation progresses ^41^. High-throughput sequencing using FAIRE-Seq, a method that identifies open and closed chromatin based on phenol separation ^42^, has revealed that, in cells that have undergone replicative senescence, previously heterochromatic domains enriched for various repeat elements become more accessible whilst euchromatic domains undergo condensation ^43^. However, it remains unknown how chromatin structure and accessibility is altered in RIS and NIS cells.

Here, we characterise the chromatin phenotype in RIS and NIS cells. We demonstrate that these two types of senescent cells exhibit distinct chromatin structures at a microscopic and nucleosome scale. Both gain multiple accessible regions or ‘nucleosome free regions’ (NFRs), which are often exclusive between RIS and NIS and reflect the transcriptional landscape of each phenotype. Strikingly, we find that autonomous and non-cell autonomous activation of the NOTCH signalling pathway in RIS cells can repress SAHFs and the formation of RIS-specific NFRs, partially by transcriptional repression of HMGAs. Our study demonstrates that chromatin structure and the nucleosome landscape can be regulated through juxtacrine signalling. The relationship between these two prominent tumour associated genes, HMGA and NOTCH1, may also have prognostic value *in vivo*.

## RESULTS

### NOTCH1 reprograms chromatin structure and abrogates SAHFs

We have previously demonstrated that ectopic NOTCH1 intracellular domain (N1ICD), an active form of NOTCH1 (Fig. 1a), can drive NIS that is distinct from RIS in terms of SASP composition ^15^. We noticed that NIS cells also have a unique chromatin structure and sought to investigate how this compares to RIS.

**Figure 1.**
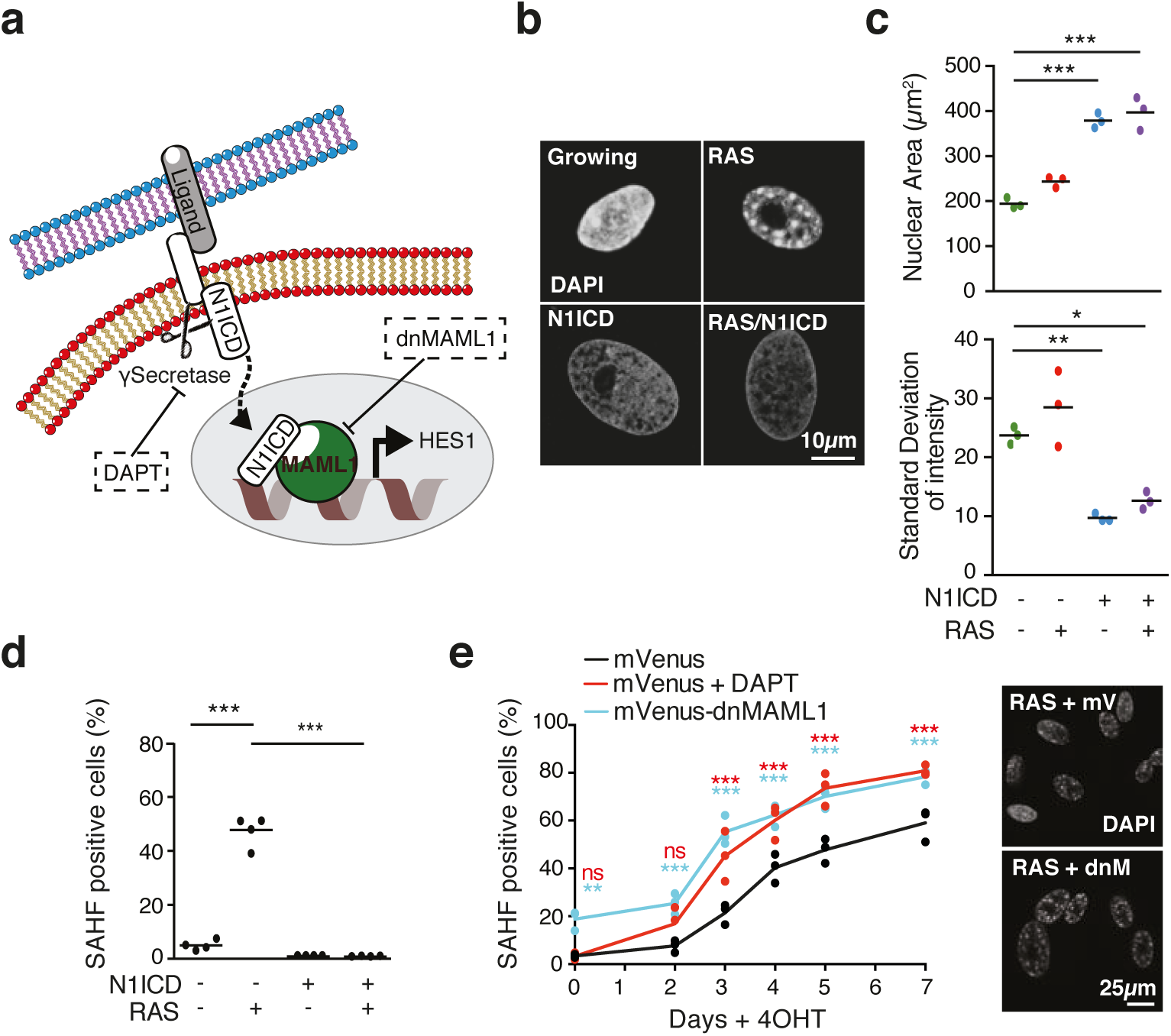
NOTCH1 signalling has a chromatin ‘smoothening’ effect that strongly blocks SAHFs in the context of RAS-induced senescence. **(a)** Diagram illustrating the NOTCH1 signalling pathway, which can be repressed chemically using DAPT or genetically by expressing dominant-negative MAML1 (dnMAML1). **(b)** IMR90 ER:HRAS^G12V^ cells infected with control vector or FLAG-N1ICD ± 4OHT for 6 days. Representative images of nuclei stained with DAPI for the conditions indicated. **(c, d)** Quantification of nuclear area, standard deviation of DAPI intensity **(c)** and number of SAHF positive cells **(d)** for the conditions indicated as in b. Lines indicate the mean value of individual replicates. n=3 **(c)** and n=4 **(d)** biologically independent replicates. **(e)** Time series analysis of SAHF positive nuclei following the addition of 4OHT to IMR90 ER:HRAS^G12V^ cells in the presence or absence of ectopic dnMAML1 or DAPT treatment. n=3 biologically independent replicates. Statistical significance calculated using one-way ANOVA with Tukey’s correction for multiple comparisons. *P ≤ 0.05, **P ≤ 0.01, ***P ≤ 0.001.

To examine the relationship between NOTCH1 and chromatin structure, we introduced ectopic N1ICD into IMR90 human diploid fibroblasts (HDFs) stably expressing a 4-hydroxytamoxifen (4OHT) inducible estrogen receptor - oncogenic HRAS fusion protein (IMR90 ER:HRAS^G12V^ cells) ^44^. Ectopic expression of N1ICD alone induced senescence with dramatically enlarged nuclei, even larger than in RIS (Fig. 1b, c). Similarly to RIS, NIS cells formed SADS, a more common chromatin feature of senescence than SAHFs (Supplemental Fig. 1a) ^26^. However, in marked contrast to RIS, where cells develop prominent SAHFs, NIS cells lacked SAHFs (Fig. 1b, d).

To ask whether NIS cells simply lack SAHFs or whether N1ICD actively modulates chromatin structure, we expressed N1ICD in the presence of HRAS^G12V^. Interestingly, N1ICD in the context of RIS also resulted in a dramatic enlargement of nuclei but a complete ablation of SAHF formation (Fig 1b-d). This was emphasised by a ‘smoothening’ of chromatin as indicated by a marked reduction in the standard deviation of DAPI signal measured within individual nuclei (Fig. 1b, c; Supplemental Fig. 1b-d), which was apparent even in the presence of HRAS^G12V^ induction. We have previously shown that ectopic N1ICD in the RIS context results in senescence with distinct SASP components largely similar to NIS ^15^. Thus, our data indicate that NOTCH is dominant over RIS in terms of chromatin phenotype as well as SASP composition, and that NOTCH signalling inhibits chromatin condensation including SAHF formation.

In IMR90 ER:HRAS^G12V^ cells, RIS develops progressively over a time period of ~ 6 days following the addition of 4OHT ^44^. NOTCH1 signalling is temporally regulated during RIS, where cleaved and active N1ICD is transiently upregulated before downregulation at full senescence ^15^. To examine temporal effects of NOTCH1 signalling on SAHF formation, we performed a time course experiment in IMR90 ER:HRAS^G12V^ cells. Cells were retrovirally infected with a dominant negative form of MAML1 fused to mVenus (dnMAML1-mVenus) or treated with the γ-secretase inhibitor DAPT to repress downstream signalling by N1ICD (Fig. 1a). We found that a greater number of SAHF-positive cells were formed and that these accumulated at earlier time points when NOTCH1 signalling was repressed (Fig. 1e). Furthermore, a dose dependent effect was evident where higher concentrations of DAPT resulted in a greater proportion of cells developing SAHF during RIS (Supplemental Fig. 1e). SAHFs are not typically prominent in DNA damage-induced senescence (DDIS) in IMR90 cells ^45^. However, DAPT significantly promoted SAHF formation in DDIS (by etoposide) (Supplemental Fig. 1f). Together, our data suggest that NOTCH signalling has a chromatin ‘smoothing’ effect that abrogates SAHF formation.

### Non-cell autonomous regulation of SAHFs

N1ICD-expressing cells can induce NIS in adjacent normal cells, at least in the case of IMR90 cells ^15^. To determine whether N1ICD-expressing cells can also alter chromatin structure in adjacent cells, we performed co-cultures between mRFP1-expressing IMR90 ER:HRAS^G12V^ cells and IMR90 cells retrovirally infected with doxycycline (DOX)-inducible FLAG-N1ICD (iN1ICD) in the presence and absence of 4OHT and DOX (Fig. 2a). Strikingly, co-culture with N1ICD-expressing IMR90 cells was sufficient to repress SAHF formation in adjacent RIS (red) cells (Fig 2b, c).

**Figure 2.**
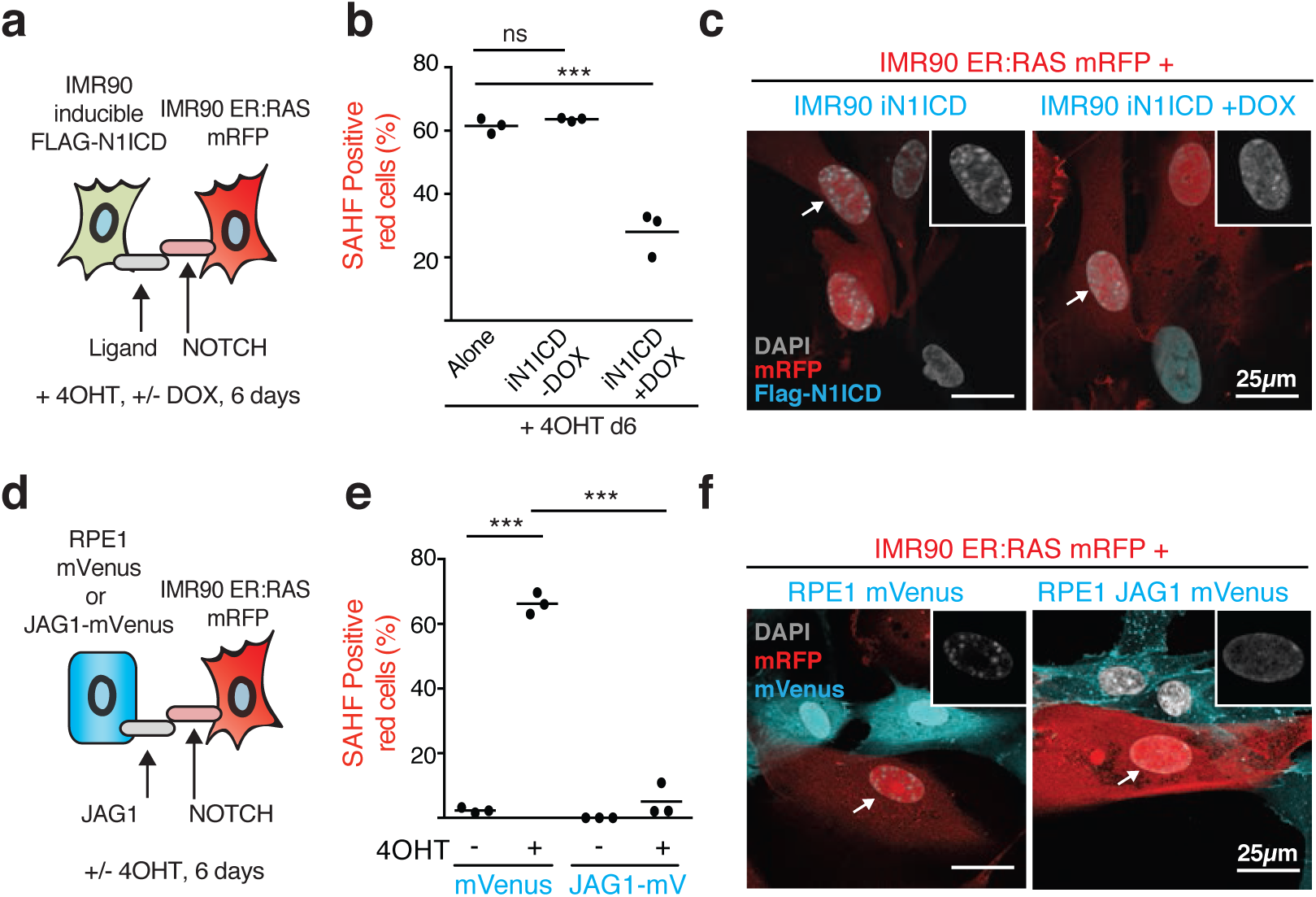
NOTCH1 signalling and JAG1 expression can non-autonomously repress SAHFs in adjacent cells. **(a)** Schematic showing experimental setup. IMR90 cells expressing either Doxycycline (DOX)-inducible FLAG-N1ICD or ER:RAS were co-cultured on 4OHT for 6 days with/without DOX. **(b)** Quantification of SAHF positive red cells for the experiment outlined in a. Alone, mono-cultured IMR90 ER:HRAS^G12V^ cells; iN1ICD, DOX-inducible N1ICD. **(c)** Representative images of co-cultures indicated. Insets are unmerged DAPI images of indicated cells (arrows) **(d)** Schematic showing experimental setup. IMR90 ER:HRAS^G12V^ cells were co-cultured with RPE1 cells stably expressing either mVenus or JAG1-mVenus for 6 days with/without 4OHT. **(e)** Quantification of SAHF positive red cells for the experiment outlined in d. **(f)** Representative images of co-cultures indicated. Insets are unmerged DAPI images of indicated cells (arrows) **(b, c)**. Lines indicate the mean value of individual replicates. n=3 biologically independent replicates for all conditions; Statistical significance calculated using one-way ANOVA with Tukey’s correction for multiple comparisons; *P ≤ 0.05, **P ≤ 0.01, *P ≤ 0.001.

Of the canonical NOTCH1 ligands, we have previously observed a strong and unique upregulation of JAGGED1 (JAG1) following ectopic N1ICD expression, which we found to be responsible for the juxtacrine transmission of senescence ^15^. We reasoned that N1ICD-mediated upregulation of JAG1 and subsequent ‘lateral induction’ of NOTCH1 signalling is a likely mechanism by which SAHFs are regulated non-autonomously. To test this hypothesis we expressed ectopic JAG1 fused to mVenus (JAG1-mVenus) in hTERT-immortalised human Retinal Pigment Epithelial (RPE1) cells. We confirmed cell surface expression of ectopic JAG1 by flow-cytometry (Supplemental Fig. 2a) before co-culturing with IMR90 ER:HRAS^G12V^ mRFP1 cells. Whilst control RPE1 cells had no effect on SAHF formation in red IMR90 RIS cells, RPE1 JAG1-mVenus cells significantly repressed the formation of SAHFs (Fig 2e, f). Note this repression did not occur when these two types of cells were co-cultured without any physical contact in a transwell format (Supplemental Fig. 2b). Our data suggest a mechanism by which lateral induction of NOTCH signalling by JAG1 can block SAHFs in the context of RIS and demonstrate a novel mechanism by which higher-order chromatin structure can be regulated through cell-cell contact.

### NOTCH signalling inhibits HMGA expression

To unravel the mechanisms underpinning NOTCH1 dependent repression of SAHFs, we re-analysed previously published RNA-seq data generated from IMR90 cells expressing HRAS^G12V^ and N1ICD ^15^. We observed that N1ICD dramatically represses the expression of *High-Mobility Group AT-hook 1* (*HMGA1*) and *HMGA2* (Supplemental Fig. 3a), critical components of SAHF structure ^27^.

To further validate that NOTCH1 signalling represses HMGA expression, we introduced constitutive N1ICD into ER:HRAS^G12V^ expressing IMR90 cells. Ectopic N1ICD significantly repressed HMGA1 and HMGA2 at an mRNA and protein level in both the presence and absence of 4OHT-induced HRAS^G12V^ (Fig. 3a, b). N1ICD has a similar effect on HMGA expression when expressed in other cell lines in the absence of HRAS^G12V^, suggesting a conserved mechanism (Supplemental Fig. 3b). In the DOX-inducible FLAG-N1ICD system, inhibition of NOTCH1 signalling by co-expression of dnMAML1-mVenus was sufficient to rescue N1ICD-mediated repression of HMGA1/ 2 (Fig. 3c, d), suggesting the effect is dependent on the canonical pathway of NOTCH signalling.

**Figure 3.**
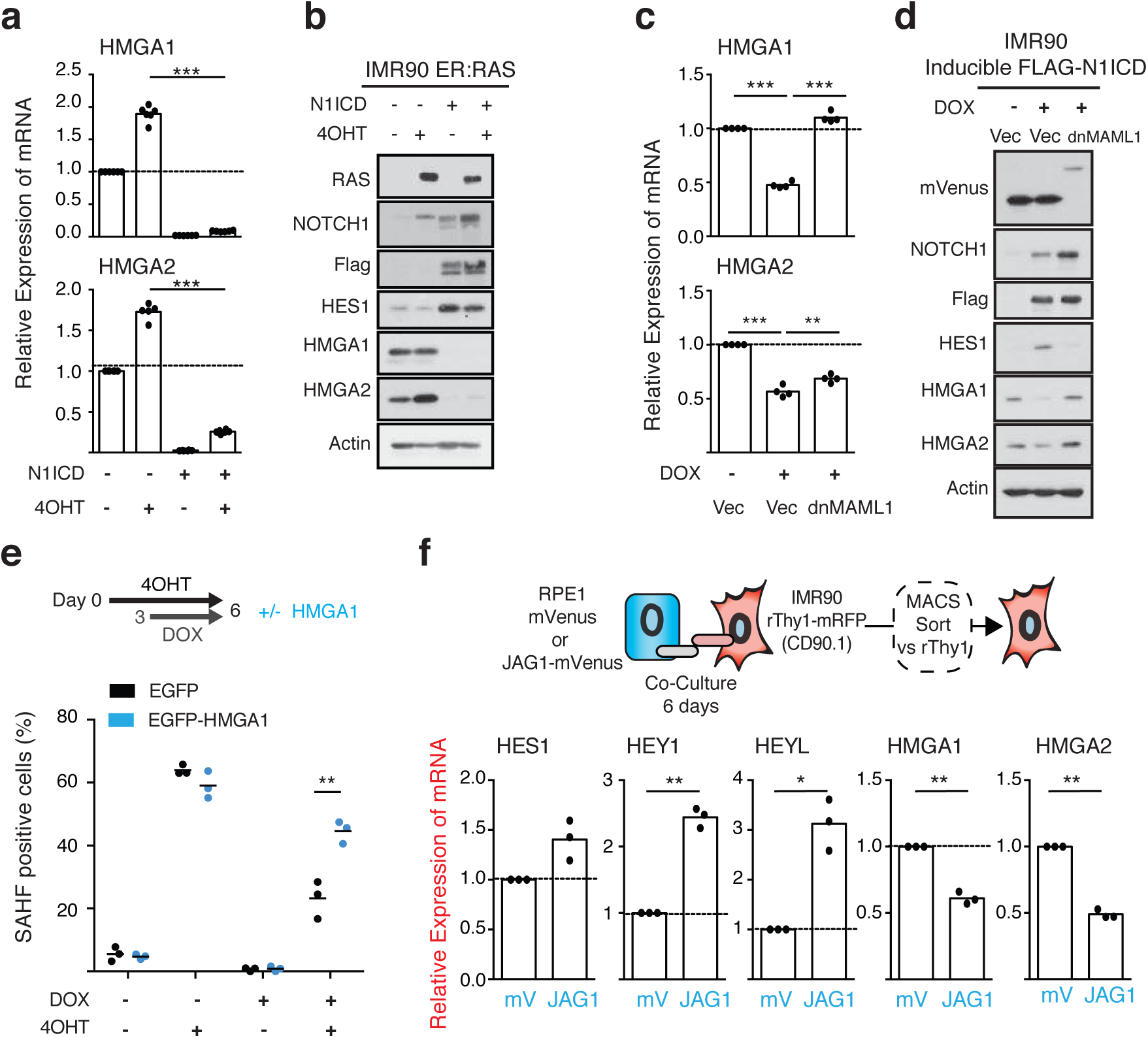
NOTCH1 signalling represses SAHFs partially by repressing HMGA proteins. **(a, b)** qRT-PCR (n=6) **(a)** and immunoblotting **(b)** for the indicated mRNA and proteins in IMR90 ER:HRAS^G12V^ cells stably infected with control vector or FLAG-N1ICD ± 4OHT for 6 days. **(c, d)** qRT-PCR (n=4) **(c)** and immunoblotting **(d)** of IMR90 cells expressing doxycycline (DOX)-inducible FLAG-N1ICD (iN1ICD) and infected with a control vector or dnMAML1 ± DOX for 3 days. **(e)** Quantification of SAHFs in IMR90 cells expressing ER:HRAS^G12V^, iN1ICD, and EGFP or EGFP-HMGA1 fusion ± 4OHT for 6 days and ± DOX for 3 days. **(f)** qRT-PCR for the indicated mRNA in IMR90 cells co-cultured with RPE1 mVenus or JAG1-mVenus cells and sorted using MACS (n=3). Statistical significance calculated using one-way ANOVA with Tukey’s correction for multiple comparisons **(a, c)** or paired t-test **(e, f)**. *P ≤ 0.05, **P ≤ 0.01, ***P ≤ 0.001.

Finally, we used IMR90 ER:HRAS^G12V^ cells expressing DOX-inducible FLAG-N1ICD to investigate whether ectopic re-expression of EGFP-tagged HMGA1 is sufficient to rescue SAHFs. The introduction of EGFP-HMGA1 resulted in a partial but significant rescue of SAHF-positive cells when cells were treated with DOX and 4OHT (Fig. 3e).

Collectively, our data suggest that NOTCH1 signalling represses the formation of SAHFs at least partially by inhibiting HMGAs, establishing a novel connection between these two prominent cancer-associated genes.

### Non-cell autonomous inhibition of HMGA

To determine whether HMGAs are repressed non-autonomously by JAG1 expressing cells, we performed further co-cultures between RPE1 cells retrovirally infected with JAG1-mVenus and IMR90 cells ectopically expressing a cell surface marker, rat Thy1, allowing for subsequent isolation using magnetic-activated cell sorting (MACS) (Fig. 3f). As expected, IMR90 cells co-cultured with JAG1 expressing cells expressed canonical NOTCH1 target genes such as *HEY1* and *HEY-L*. Strikingly, both *HMGA1* and *HMGA2* were significantly repressed in IMR90 cells cultured with JAG1 expressing RPE1 cells when compared to controls (Fig. 3f), demonstrating that HMGA proteins can be repressed non-autonomously.

### Gene-distal nucleosome free regions accumulate in RIS and NIS

To investigate whether NOTCH1 influences chromatin structure at a higher resolution, we employed ATAC-seq (Assay for Transposase-Accessible Chromatin using Sequencing) ^46^. This method takes advantage of a hyperactive Tn5 transposase enzyme that inserts sequencing adapters into accessible chromatin, or ‘nucleosome-free regions’ (NFRs). Following PCR amplification these regions can be sequenced as a method of identifying NFRs genome wide (Fig. 4a). Indeed, chromatin accessibility is known to be a direct reflection of cell phenotype and the transcriptional landscape ^40^.

**Figure 4.**
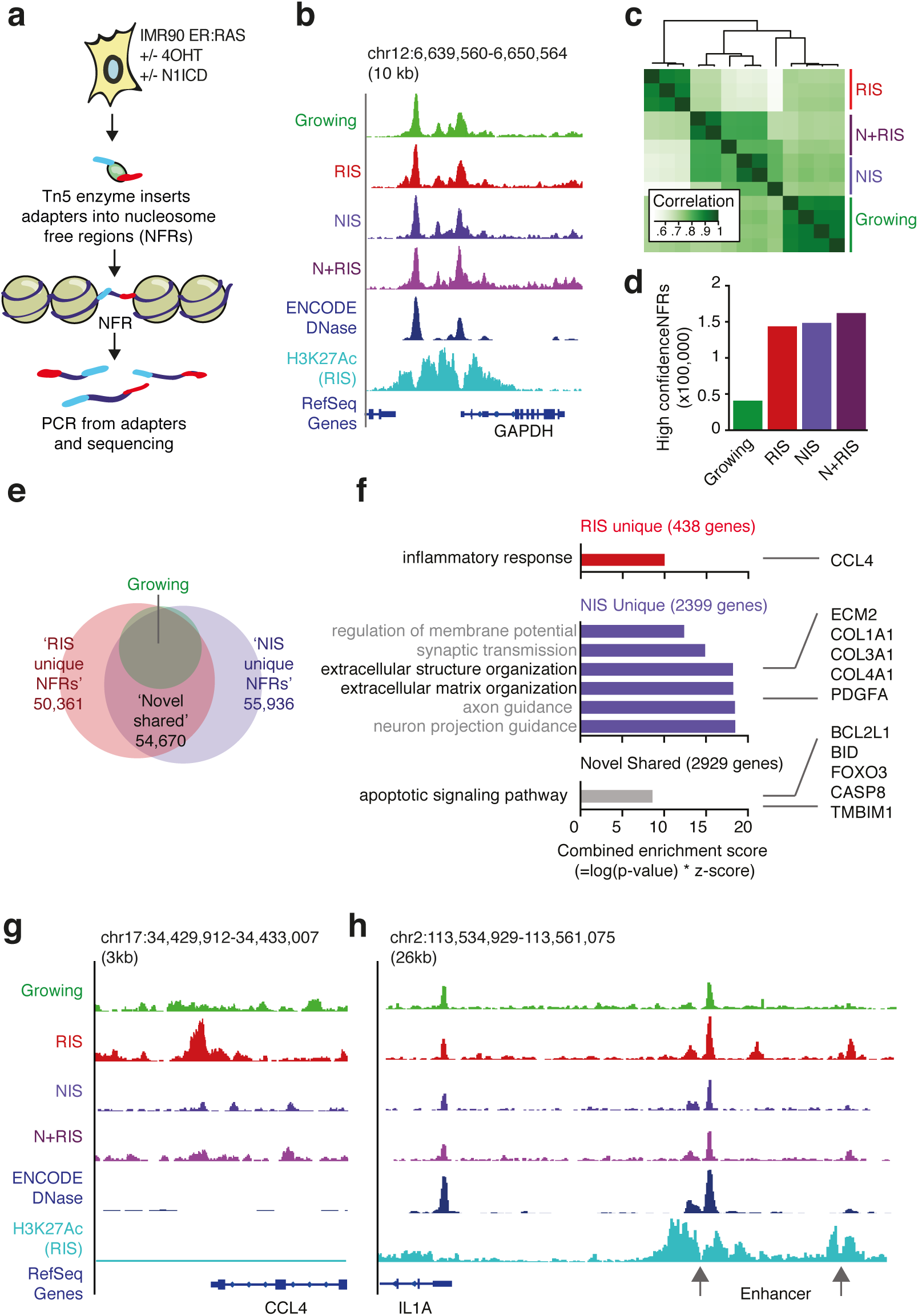
Nucleosome free regions (NFRs) reflect gene transcription in RIS and NIS cells. **(a)** Diagram illustrating the method, ATAC-seq. **(b)** Genome browser image showing normalised ATAC-seq coverage and the enhancer associated histone modification, H3K27Ac, in the cell conditions indicated around the GAPDH locus. DNase I sensitive regions in normal human lung fibroblast (NHLF) cells from ENCODE is shown for comparison. **(c)** Unbiased clustering of replicates for the cell conditions indicated. **(d)** Number of high-confidence NFRs (present in at least two replicates) for the cell conditions indicated. **(e)** Venn-diagrams showing overlap between NFRs identified in the conditions indicated. **(f)** Gene ontology analysis (GO biological process 2015) using promoter proximal NFRs identified in e, with genes of interest in each category indicated. **(g, h)** Representative genome browser images showing normalised ATAC-seq coverage files and H3K27Ac around two regions that are substantially altered between cell conditions.

We generated at least 3 replicates from IMR90 ER:HRAS^G12^ cells expressing FLAG-N1ICD or a control vector and induced with 4OHT or not. For simplicity, these conditions were labelled as ‘Growing’, ‘RIS’, ‘NIS’ and ‘N+RIS (expressing both N1ICD and RAS)’. Using a previously published trimmed-means normalisation approach ^47^, we first generated normalised coverage files. When interrogated using a genome browser, these were comparable to each other and to ENCODE DNase-sequencing data from normal human lung fibroblasts (NHLF), especially around housekeeping genes (Fig. 4b). Most of the samples, excluding a single replicate from the NIS and N+RIS conditions, were of high quality with a reads in peaks percentage (RiP%) of more than 10% (Supplemental Fig. 4a). Replicates clustered well by unbiased clustering and PCA analyses (Fig. 4c, Supplemental Fig. 4b).

To generate a consensus set of high confidence NFRs for downstream analyses, we identified peaks present in at least two replicates of each condition (Fig. 4d, Supplemental Table 1). We detected 40,903 peaks in growing IMR90 cells, 143,827 peaks in RIS cells and 147,468 peaks in NIS cells. Therefore, a large number of novel NFRs were gained in RIS (104,579) and NIS (108,839) cells when compared to growing IMR90 cells. In contrast, only a small number of the NFRs present in growing cells were lost in RIS (1,655) and NIS (2,274) cells.

A previous study mapping chromatin accessibility in replicatively senescent cells using FAIRE-Seq found that gene-distal regions, especially repeat regions, become relatively more open whilst genic regions become closed when compared to growing fibroblasts ^43^. Consistently, NFRs in RIS and NIS cells (when compared to growing cells) were enriched at gene distal sites with the majority of new peaks mapping to enhancer, intergenic, intronic and repeat regions (Supplementary Fig. 4c, d). Many of these repeat regions were further annotated as LINEs, LTRs, SINEs, Satellites and Simple repeat regions (Supplementary Fig. 4e). In contrast to replicative senescence described in the previous study ^43^, we did not observe a reduction in the number of NFR’s mapping to genic regions in RIS and NIS cells, though the increase at transcriptional start sites (TSS), exons and CpG islands was less dramatic (Supplementary Fig. 4d). Therefore, unlike in replicative senescence, RIS and NIS may be characterised by global chromatin opening at the nucleosome scale.

### NFRs in RIS and NIS reflect gene transcription

Gene promoters of transcribed genes must first be accessible to transcription factors and DNA-polymerase; therefore ATAC-seq data should correlate well with gene transcription ^39,40^. To determine whether NFRs in RIS and NIS reflect gene transcription, we first intersected consensus peaks detected in growing, RIS and NIS cells (Fig. 4e, Supplemental Table 2). We isolated 50,361 peaks unique to RIS cells (‘RIS unique NFRs’), 55,936 peaks unique to NIS cells (‘NIS unique NFRs’) and 54,670 peaks present in both RIS and NIS cells that were not detected in growing cells (‘Novel shared NFRs’). All three of these subsets had a tendency to be gene distal when compared to peaks found in growing IMR90 cells, with RIS-unique NFRs showing the greatest tendency (Supplemental Fig. 5a).

Next, we mapped NFRs within 300bp of a gene TSS to genes (in order to confidently map NFRs to genes) and performed Gene Ontology (GO) analyses with these lists. Consistent with our previous RNA-seq data ^15^, genes with RIS-unique NFRs were significantly enriched in the term ‘inflammatory response’ whilst genes with NIS-unique NFRs were enriched in the terms ‘extracellular structure organization’ and ‘extracellular matrix organization’ and included genes that transcribe fibrotic proteins such as collagens (Fig. 4f).

Genes with ‘Novel shared NFRs’ were enriched in the term ‘apoptotic signalling pathway’, which was particularly striking as both pro- and anti-apoptotic proteins have recently been described to play important roles in senescence (Fig. 4f) ^48-50^. For example, we detected a novel shared NFR at the promoter of the gene *BCL2L1*, which encodes the protein BCL-XL, an anti-apoptotic protein and potential target for ‘senolytic drugs’ ^49,50^. The increased accessibility at the promoters of these genes in both RIS and NIS suggests a unique mode of apoptotic sensitivity in senescent cells, which are generally apoptosis-resistant but can be selectively killed using senolytics.

Finally, we asked whether the genes in our gene lists are differentially expressed in RIS and NIS cells by RNA-seq. To do so, we reanalysed previously generated data ^15^ and found that, on average, genes with RIS-specific NFRs were transcriptionally upregulated in RIS cells, whilst genes with NIS-specific NFRs were transcriptionally upregulated in NIS cells (Supplemental Fig. 5b, Supplemental Table 3). Normalised ATAC-seq coverage files, when viewed using a genome browser (Fig. 4g, h, Supplemental Fig. 5c), supported these observations. We also noted that, whilst few inflammatory genes gained RIS-unique NFRs at their promoters, some had upstream enhancer elements that become accessible in RIS cells (Fig. 4g, Supplemental Fig. 5d). These data demonstrate that RIS and NIS cells have unique open chromatin landscapes reflecting their specific transcriptional profiles.

### NOTCH signalling antagonises the formation of RIS specific NFRs

By unbiased clustering, we observed a greater correlation between NIS and N+RIS cells than between RIS and N+RIS cells (Fig 4c, Supplemental Fig. 4b). This suggests a dominant effect of N1ICD over RAS on the nucleosome scale, which is consistent with our previous observations for SASP components ^15^ and SAHFs (Fig. 1d).

To validate whether NFRs in RIS are repressed by N1ICD, we took the 50,361 ‘RIS-unique NFRs’ (Fig. 4e) and intersected them with the consensus peaks identified in the N+RIS condition. Of the RIS unique NFRs, 61% (30,775) were repressed by N1ICD (Fig. 5a). HMGA proteins have previously been shown to affect chromatin compaction by competing with Histone-H1 for linker DNA. To determine whether repression of HMGA1 is a mechanism by which N1ICD can repress RIS-unique NFRs, we generated additional ATAC-seq samples from IMR90 ER:HRAS^G12^ cells expressing a short-hairpin against HMGA1 ^27^ and treated with 4OHT, hereafter referred to as ‘RIS+shHMGA1’. By intersecting the 50,361 RIS-unique NFRs with consensus peaks identified in the RIS+shHMGA1 condition, we found that 30% (15,292) were dependent on HMGA1 (Fig. 5a). Of these, 77.6% (11,879) were also repressed by N1ICD. These analyses illustrate that a subset of RIS-unique NFRs can be repressed by N1ICD, possibly through HMGA downregulation. RIS-unique NFRs were significantly more AT-rich than other subsets of NFRs (Fig. 5b), further supporting the involvement of HMGA in formation of RIS-unique NFRs. These data suggest that, in the RIS context, HMGA1 is a key regulator of chromatin structure at the nucleosome and microscopic level.

**Figure 5.**
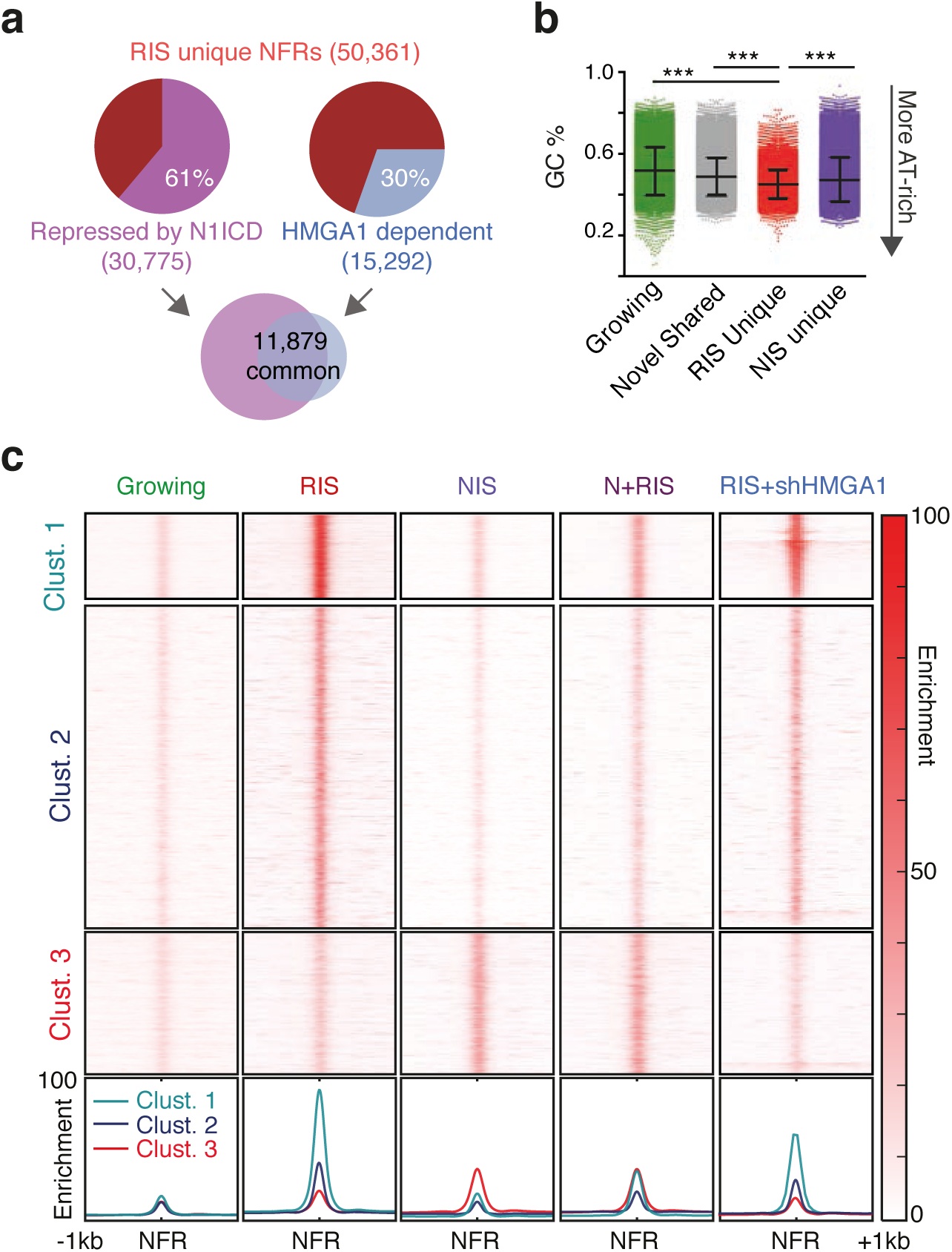
Ectopic N1ICD and HMGA1 knockdown antagonise formation of RIS specific NFRs. **(a)** Number of ‘RIS specific NFRs’ identified in Fig. 4e that are repressed by N1ICD (not detected in RIS+N1ICD ATAC-seq samples) and are HMGA1 dependent (not detected in RIS+shHMGA1 ATAC-seq samples) and the intersect between these two subsets. **(b)** Genomic GC percentage of NFRs in the subsets indicated, taken from Fig. 4e. Statistical significance calculated using one-way ANOVA with Tukey’s correction for multiple comparisons; *P ≤ 0.05, **P ≤ 0.01, ***P ≤ 0.001. **(c)** Heat-map showing the enrichment of reads from normalised coverage files around NFRs detected in any cell condition other than in growing IMR90 cells (top) and summary plots showing mean enrichment (bottom).

To validate the above ‘binary’ approach, where we assume that peaks are either present or absent, we used our normalised coverage files to perform unbiased *k-means* clustering around novel consensus NFRs identified in RIS, NIS or N+RIS conditions (that were detected in growing cells) (Fig. 5c). NFRs separated into 3 clusters: clusters 1 and 2 were dominated by RIS whilst cluster 3 was dominated by NIS. Clusters 1 and 2 appeared to separate largely due to the level of enrichment, where NFRs in cluster 1 were stronger. Strikingly, the signal in clusters 1 and 2 was reduced in the N+RIS and RIS+shHMGA1 conditions when compared to the RIS condition (Fig. 5c). Notably, whilst peaks in cluster 3 were increased in the N+RIS condition, they did not increase in the RIS+shHMGA1 condition, suggesting an HMGA-independent mechanism of chromatin opening in NIS (Fig. 5c). Therefore, in line with microscopic SAHF structures, N1ICD alters chromatin structure in RIS at the nucleosome scale in part through repressing HMGA expression.

### Non-cell autonomous regulation of chromatin by tumour cells

HMGA1 and NOTCH1 are two prominent cancer-associated genes. Both can act as oncogenes or tumour suppressors in a context-dependent manner. We reasoned that the relationship between these two genes might also be important in the tumour microenvironment and asked whether tumour cells expressing JAG1 can affect HMGA expression and chromatin structure in adjacent fibroblasts.

To answer this question, we used the Cancer Cell Line Encyclopaedia (CCLE) ^51^ to identify tumour cell lines that express low (MCF7), medium (A549) and high (Hep3B) levels of JAG1, which we confirmed by immunoblotting (Fig. 6a). Co-culture of tumour cell lines with ER:HRAS^G12V^ and mRFP1 expressing IMR90 cells in the presence of 4OHT was sufficient to repress SAHF formation in red RIS cells in a contact-dependent manner (Fig. 6b; Supplemental Fig. 6a, b). The number of SAHF-positive red cells correlated with the level of JAG1 expressed by the tumour cell lines (Fig. 6b). Inhibition of SAHF formation in the co-culture was completely abrogated by DAPT, suggesting the effect is dependent on the canonical NOTCH pathway (Fig. 6b).

**Figure 6.**
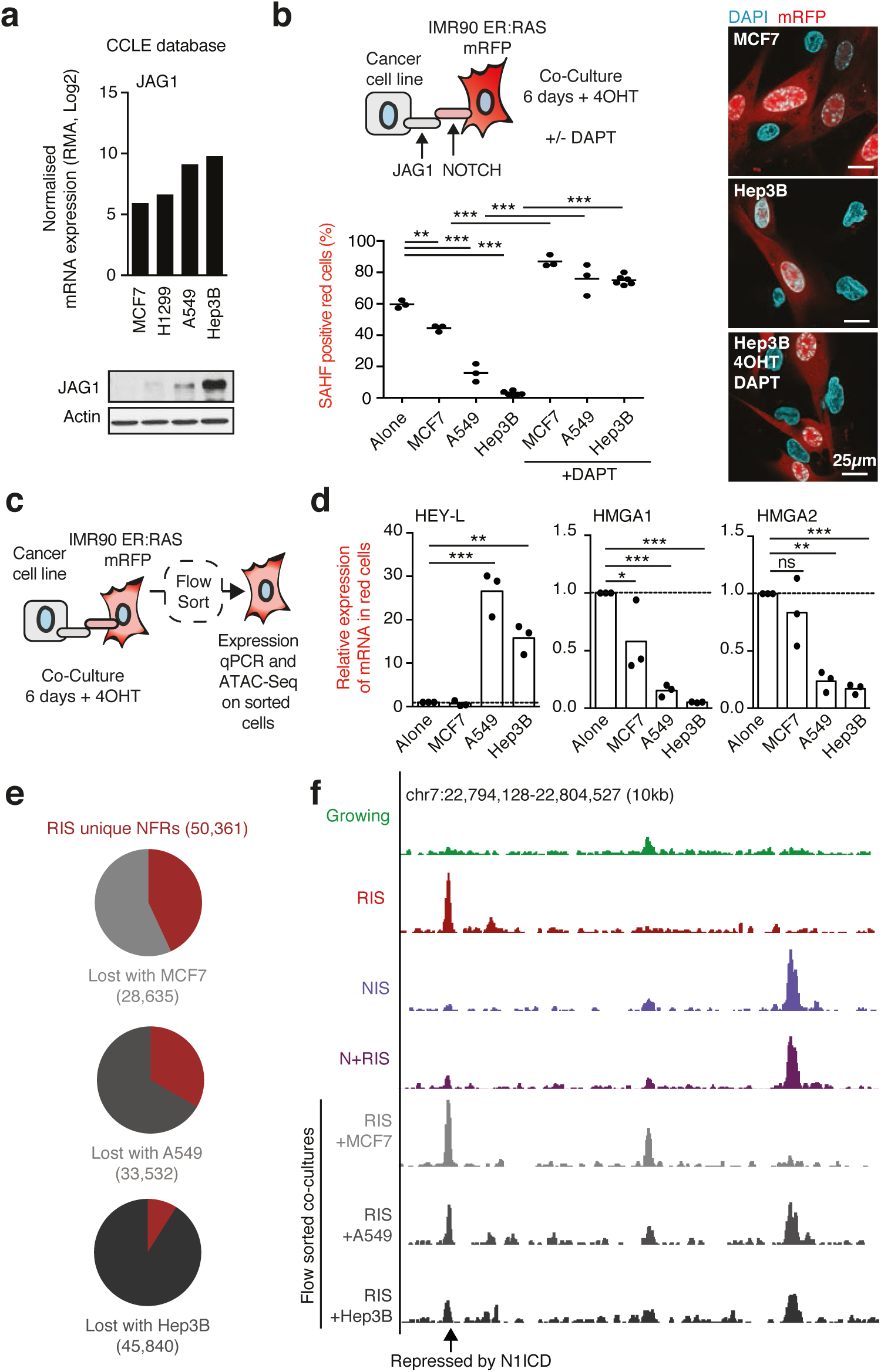
Tumour cells can repress SAHFs and RIS specific NFRs in adjacent RIS fibroblasts. **(a)** Normalised mRNA expression values from the Cancer Cell Line Encyclopaedia (CCLE) and immunoblotting of JAG1 in the mour cell lines indicated. **(b)** Quantification of SAHFs in red cells of co-culture between IMR90 ER:HRAS^G12V^ RFP1 cells and tumour cell lines indicated for 6 days + 4OHT ± DAPT. **(c, d)** Experimental setup **(c)** and qRT-PCR **(d)** of mRNA isolated from flow sorted IMR90 ER:HRAS^G12V^ mRFP1 cells cultured with tumour cell lines OHT for 6 days relative to cells cultured alone. n=3 biologically independent replicates. Statistical significance lculated using one-way ANOVA with Tukey’s correction for multiple comparisons; *P ≤ 0.05, **P ≤ 0.01, ***P ≤ **(e)** Number of ‘RIS specific NFRs’, identified in Fig. 4e, that are lost when IMR90 ER:HRAS^G12V^ mRFP1 cells are cultured with the tumour cell lines indicated before flow sorting as in described in c. **(f)** Representative genome owser images showing normalised ATAC-seq coverage files around an altered region in the cell conditions indicated and described in e. Note that ‘+MCF7’, ‘+A549’ and ‘+Hep3B’ denotes that RIS cells have been previously co-cultured with these cell lines for 6 days prior to flow sorting.

To determine whether tumour cell lines can induce NOTCH1 signalling and repress HMGAs non-autonomously, we repeated the co-cultures and isolated the IMR90 ER:HRAS^G12V^ mRFP1 cells using flow cytometry (Fig. 6c). Quantitative RT-PCR showed a dramatic upregulation of the canonical NOTCH1 target gene *HEYL* and a concurrent downregulation of *HMGAs* in fibroblasts co-cultured with JAG1-expressing tumour cells, particularly A549 and Hep3B cells (Fig 6d). Two other canonical target genes, *HES1* and *HEY1*, were not as dramatically upregulated by JAG1 expressing cell lines (Supplemental Fig. 6c), which highlights the complexity of the NOTCH pathway but may also suggest that *HEYL* plays an important role in this interaction.

Finally, we asked whether tumour cell lines could repress ‘RIS-unique NFR’ formation in fibroblast cells, as was the case for ectopic N1ICD (Fig. 5a). We isolated 4OHT-induced IMR90 ER:HRAS^G12V^ mRFP1 cells from co-culture with tumour cell lines by flow cytometry and performed ATAC-seq using these cells (Fig. 6c). By performing intersections between RIS-unique NFRs (50,361) (Fig. 4e) and consensus peak sets from red RIS cells previously co-cultured with tumour cell lines, we found that co-culture with MCF7, A549, or Hep3B cells repressed 57% (28,635), 67% (33,532) or 91% (45,840) of RIS-unique NFRs respectively (Fig. 6e, f). These data correlated well with the ability of the tumour cell lines to repress SAHFs in adjacent IMR90 (Fig. 6b). NFRs repressed by co-culture with the tumour cell lines overlapped well with each other and with the 15,292 ‘HMGA1-dependent’ NFRs (Supplemental Fig. 6d, e). These data suggest that tumour cells expressing JAG1 can dramatically alter the chromatin landscape of adjacent stromal cells in a juxtacrine manner, particularly in the senescence context where HMGA proteins are highly expressed.

### HEY-L and HMGA1 anti-correlate in multiple tumour types

If NOTCH1 signalling inhibits HMGA1 *in vivo*, we would expect an anti-correlation between NOTCH1 activity and HMGA1 expression in human tumour samples. To test whether this is the case, we first performed a pan-tissue type analysis using microarray data from the R2 database (http://r2.amc.nl) by comparing the expression of HMGA1 and canonical NOTCH1 target genes (Supplemental Fig. 7a, b, c). When Z-score expression values were compared across all the datasets in R2, we observed a strong negative correlation between *HMGA1* and *HEY-L* (R=-0.356, p < 0.0001) and between *HMGA1* and *HEY1* (R=-0.281, p <0.0001), but no correlation between *HMGA1* and *HES1* (Supplemental Fig. 7a, b, c). This result was particularly interesting, because HEY-L was also the most dramatically upregulated gene in IMR90 fibroblasts co-cultured with JAG1-expressing RPE1 cells (~3 fold) (Fig. 3g) and tumour cell lines (~20 fold) (Fig. 6d). Using the web-based tool KM plotter ^52^, which incorporates microarray data from 5,143 breast ^53^ and 2,437 lung cancer ^54^ patients, we found that patients with low *HMGA1* or high *HEY-L* have a better prognosis than patients with the inverse expression pattern in lung and breast cancer (Supplemental Fig. 8a, b), suggesting that the relationship between these proteins may have prognostic value.

As microarray data can be dependent on the quality of the probe used, we analysed the expression of *HMGA1* and *HEY-L* using RNA-seq data generated by the TCGA Research Network ^55^ (http://cancergenome.nih.gov). Consistently, there was a significant negative correlation between these two genes in the majority of tumour types analysed (Fig. 7a). There was a particularly strong anti-correlation in Lung Squamous Cell (SCC) (Fig. 7b) (r=0.4842; p= < 0.0001). Moreover, when further categorised based on expression into ‘*HEY-L* high - *HMGA1* low’ and ‘*HEY*-L low - *HMGA1* high’ tumours we found that patients in the former category had a significantly better prognosis for Lung SCC (Fig. 7c) (p=0.00316). These data demonstrate that an anti-correlation between HMGA1 and NOTCH1 activity is evident in cancer and that this correlation can be prognostic of patient outcome.

**Figure 7.**
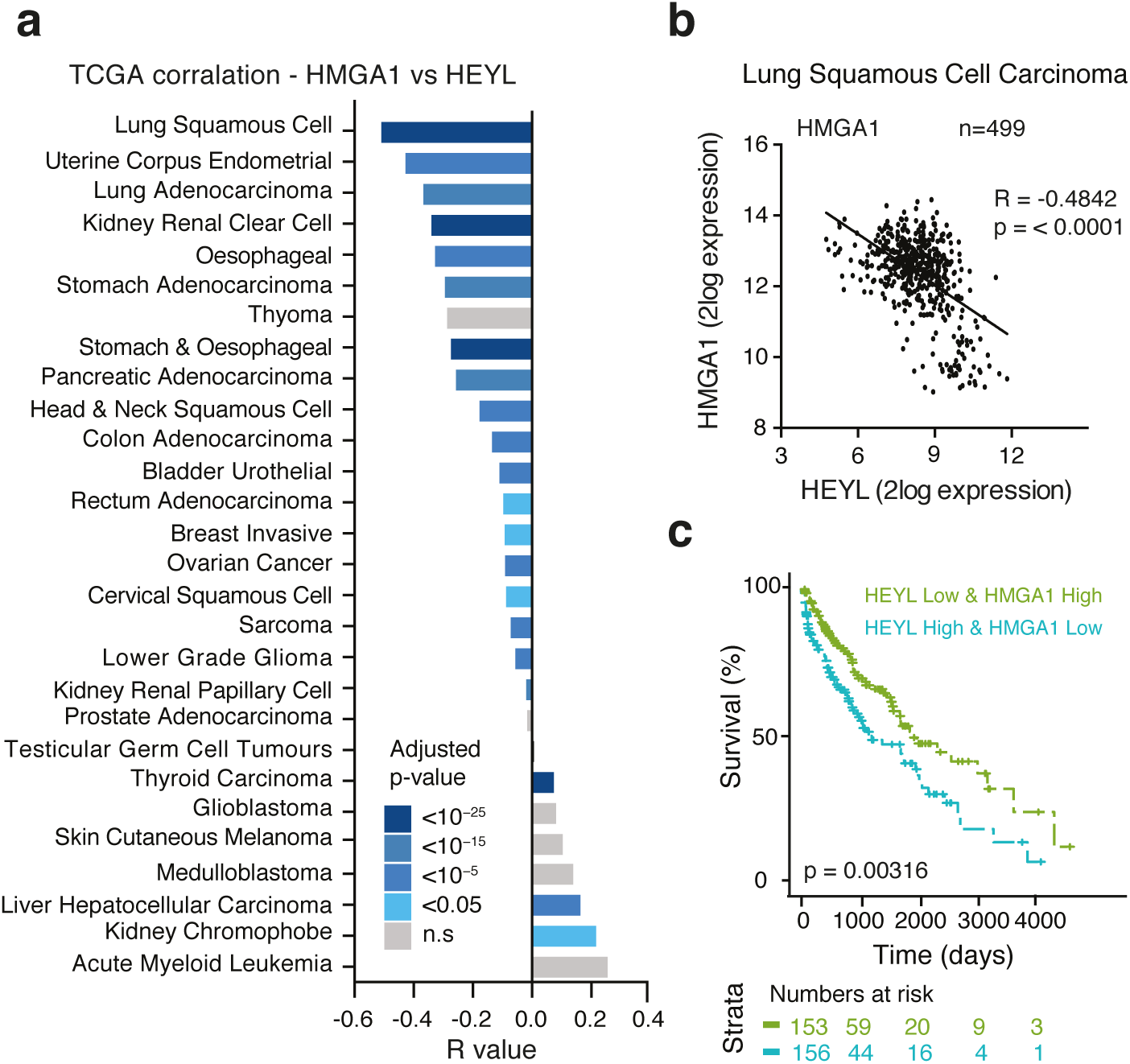
HEY-L and HMGA1 gene expression values anti-correlate in multiple tumour types. **(a)** Pan-cancer analysis of the TCGA database. Correlation between HMGA1 and HEYL is plotted in the indicated tumour types. Colours represent Bonferroni adjusted p-values on Pearson’s correlation p-values. **(b)** 2Log expression of HMGA1 against HEYL in Lung Squamous Cell Carcinoma (LSCC). **(c)** Kaplan-Meier plot showing survival of LSCC patients stratified by HMGA1 and HEYL gene expression.

## DISCUSSION

In the current study, we provide evidence for NOTCH-mediated, contact-dependent ‘lateral modulation’ of chromatin structure at both the microscopic and nucleosome scales. Whilst RIS cells form prominent SAHFs at the microscopic scale ^19,20^, at the nucleosome scale we observed a genome wide increase in chromatin accessibility, and both SAHFs and RIS-unique NFRs can be inhibited by N1ICD-mediated repression of HMGA1 (Fig. 8). While the essential and structural role for HMGA1 in SAHF formation is well established ^27^, its role in chromatin accessibility is unclear. HMGA proteins compete with Histone-H1 for linker DNA and thus affect chromatin compaction by increasing the accessibility of chromatin to other transcription factors, as demonstrated by techniques such as fluorescence recovery after photo bleaching (FRAP) and MNase digestion assays ^56-58^. Our data, using sequencing technology, demonstrate that HMGA1 facilitates the formation of ectopic NFRs. It is known that chromatin accessibility is an indicator of developmental maturity ^41^ and that cancer cells acquire ectopic NFRs ^41,59^. For example, during the metastasis of small cell lung cancer (SCLC), a dramatic increase in chromatin accessibility at distal regulatory elements allows tumour cells to co-opt pre-programmed gene expression programs, which can provide a growth advantage ^47^. Thus, our data raise an interesting possibility that HMGA1 can drive pluripotency and cancer ^36,37^ in part by modulating chromatin accessibility. It will be important to understand how HMGA1 facilitates both chromatin ‘opening’ at the nucleosome scale and the formation of SAHFs, and to determine whether the two are related. We wonder whether RIS-unique NFRs, which are mostly HMGA1-dependent and gene distal, could have structural rather than regulatory functionality.

**Figure 8.**
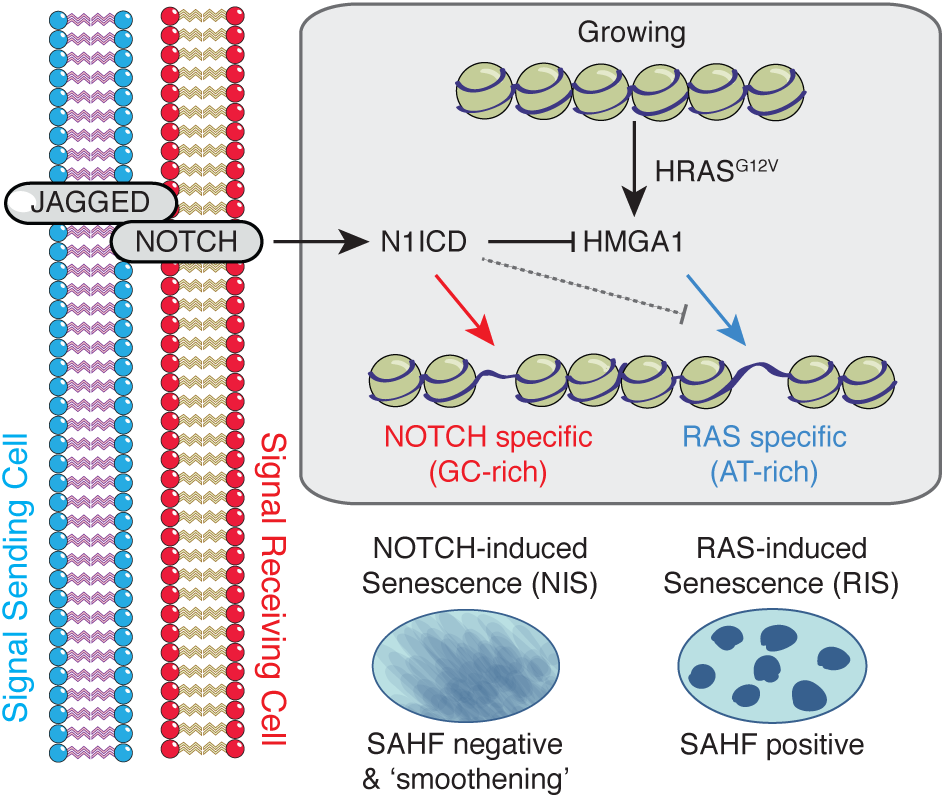
NOTCH1 signalling mediates non-cell autonomous regulation of chromatin structure at the microscopic and nucleosome scale. Lateral induction of NOTCH1 activity in a signal-receiving cell by JAG1 on the surface of an adjacent cell (including cancer cells) can drive NIS. NIS cells form ‘NIS unique NFRs’, which tend to be relatively GC rich, and microscopically ‘smoothened’ chromatin. In the context of RIS, non-cell autonomous activation of NOTCH1 signalling can repress the formation of AT-rich ‘RIS unique NFRs’ at the nucleosome level and SAHFs at the microscopic level. Mechanistically, N1ICD represses HMGA1, which is responsible for SAHF formation and at least partially for the formation of ‘RIS-unique NFRs’.

Chromatin accessibility was also increased in NIS cells although these are often at distinct loci. Unlike RIS cells, NIS cells do not form SAHFs and are instead characterised by a chromatin ‘smoothening’ phenotype. The mechanisms of chromatin smoothening in NIS and formation of NIS-unique NFRs, and whether these events are related with each other, remain unclear. Note, although knockdown of HMGA1 blocked formation of SAHFs and many RIS-specific NFRs, it was not sufficient to induce NIS-like chromatin smoothening or NIS-unique NFRs, thus NOTCH signalling modulates chromatin by both HMGA-dependent and HMGA-independent mechanisms (Fig. 5C). It is known that chromatin accessibility can be facilitated by histone acetylation ^60-63^. N1ICD activates gene transcription by recruiting histone acetyl-transferases (HATs) ^16,64,65^ and has more recently been shown to drive rapid and widespread deposition of H3K56Ac ^66^, which is know to be associated with nucleosome assembly, particularly in DNA replication or repair ^67-69^.

Our data also suggest a novel mechanism for non-cell autonomous regulation of chromatin structure. N1ICD-expressing cells can induce NIS in adjacent cells by 'lateral induction’ ^15^ and here we show that they also reprogram the chromatin landscape in adjacent cells. Considering that NOTCH1 is involved in development, it may not be surprising that it can have such a dramatic effect on chromatin structure at the nucleosome scale. A final implication of our study may be that NOTCH signalling in tumours can alter chromatin structure in adjacent stromal cells. Using the *Pten* null mouse model of prostate cancer, Su and colleagues ^70^ demonstrated that JAG1 expression in tumour cells facilitates the formation of a ‘reactive stroma’, which plays an important role in tumour development ^71-73^. It will be important to test whether chromatin structure is altered in the stroma of such tumours and whether this is dependent on HMGA repression. In NOTCH-ligand expressing tumours, targeting chromatin-modifying enzymes in the stromal compartment may present a unique therapeutic opportunity to alter the tumour niche.

## METHODS

### Cell culture

IMR90 human diploid fibroblasts (ATCC) were cultured in DMEM/10% foetal calf serum (FCS) in a 5% O_2_ / 5% CO_2_ atmosphere. hTERT-RPE1 cells (ATCC) were grown in DMEM-F12/10% FCS in a 5% O_2_ / 5% CO_2_ atmosphere. MCF7, H1299, A549 and Hep3B cells (ATCC) were grown in DMEM/10% FCS in a 5% CO_2_ atmosphere. Cell identity was confirmed by STR genotyping. Cells were regularly tested for mycoplasma contamination and always found to be negative.

Co-cultures were set up at a cell number ratio of 1:1 and performed in DMEM/10% FCS in a 5% O_2_ / 5% CO_2_ atmosphere. For transwell experiments, IMR90 cells were plated in the bottom chamber and hTERT-RPE1 or tumour cells were plated into the top chamber of a Corning 12-well Transwell plate (CLS3460 Sigma).

The following compounds were used in cultures: 4-hydroxytamoxifen (4OHT) (Sigma); *N*-[(3,5-difluorophenyl)acetyl]-L-alanyl-2-phenyl]glycine-1,1-dimethylethyl ester (DAPT) (Sigma); etoposide (Sigma); doxycycline (DOX) (Sigma).

### Vectors

The following retroviral vectors were used: pLNCX (clontech) ER:HRAS^G12V^ ^44^; pWZL–hygro for N1ICD–FLAG (residues 1758–2556 of human NOTCH1 ^15^) and mRFP1; pLPC-puro for dnMAML1-mVenus (residues 12-74 of human MAML1) ^15^, mRFP1, rThy1-mRFP1, JAGGED1-mVenus and mVenus; pQCXIH-i for DOX-inducible N1ICD-FLAG ^15^. MSCV-puro for miR30 shHMGA1 (shHMGA1 target sequence ATGAGACGAAATGCTGATGTAT ^27^).

To generate pLPC-puro rThy1-mRFP1, we first PCR cloned mRFP into pLPC-puro (pLPC-puro-x-mRFP, where x denotes cloning sites to express mRFP-fusion proteins). The CDS of rat-Thy1 was PCR amplified from cDNA (a gift from M. de la Roche, CRUK CI, UK), removing the stop codon, before cloning into pLPC-puro-x-mRFP. To generate pLPC-JAGGED1-mVenus, the CDS of human JAGGED1 was amplified using cDNA derived from N1ICD-expressing IMR90 cells, removing the stop codon, before cloning into pLPC-puro-x-mVenus.

### Flow cytometry

Analysis of ectopic JAG1-mVenus expression was conducted by flow cytometry as previously described ^15^. Cells were stained with anti-JAG1-APC (FAB1726A, R&D systems) or isotype control antibody before analysis on a FACSCalibur flow cytometer (Becton Dickenson). Flow data were further analysed using FlowJo v10.

### Magnetic-activated cell sorting and Fluorescence-activated cell sorting

Magnetic-activated cell sorting (MACS) of rThy1-expressing cells was performed using CD90.1 microbeads (130-094-523, Miltenyl Biotec) according to manufacturers instructions. Fluorescence-activated cell sorting (FACS) was performed using an Influx (Becton Dickenson) flow cytometer.

### Fluorescence microscopy

Analysis was performed as previously described ^27^. Briefly, cells were plated onto #1.5 glass coverslips the day before fixation to achieve approximately 60% confluence. Cells were fixed in 4% (v/v) paraformaldehyde (PFA) and permeabilised with 0.2% (v/v) Triton X-100 in PBS with DAPI. Coverslips were mounted onto Superfrost Plus slides (4951, Thermo Fisher) with Vectashield Antifade mounting medium (H-1000, Vector Laboratories Ltd.). Images were obtained using a Leica TCS SP8 microscope with a HC PL APO CS2 1.4NA 100x oil objective (Leica Microsystems). At least 30 nuclei were captured per biological replicate and condition before Fiji ^74^ was used to calculate nuclear area, standard deviation and maximum intensity of DAPI signal per nucleus. SADS were visualised by DNA-Fluorescence in situ hybridisation (FISH) as previously described ^26^ using fluorescent probes that target the α-satellite repeat sequence (5’-CTTTTGATAGAGCAGTTTTGAAACACTCTTTTTGTAGAATCTGCAAGTGGATATTTGG-3’). The percentage of SAHF and SADS-positive cells was counted by scoring at least 200 cells per replicate and condition.

### Quantitative RT-qPCR

RNA was prepared using the Qiagen RNeasy plus kit (74136, Qiagen) according to the manufacturer’s instructions and reverse-transcribed to cDNA using the Applied Biosystems High-Capacity Reverse Transcription Kit (43-688-13, Thermo Fischer). Relative expression was calculated as previously described ^27^ on an Applied Biosystems Quantstudio 6 by the 2−ΔΔ^*Ct*^ method ^75^ using β-actin (ACTB) as an internal control. The following primers were used:

ACTB forward: 5’-GGACTTCGAGCAAGAGATGG-3’

ACTB reverse: 5’-AGGAAGGAAGGCTGGAAGAG-3’

HEYL forward: 5’-CTCCAAAGAATCTGTGATGCCAC-3’

HEYL reverse: 5’-CCAGGGACAATGAAAGCAAGTTC-3’

HEY1 forward: 5’-CCGCTGATAGGTTAGGTCTCATTTG-3’

HEY1 reverse: 5’-TCTTTGTGTTGCTGGGGCTG-3’

HES1 forward: 5’-ACGTGCGAGGGCGTTAATAC-3’

HES1 reverse: 5’-ATTGATCTGGGTCATGCAGTTG-3’

HMGA1 forward: 5’-GAAAAGGACGGCACTGAGAA-3’

HMGA1 reverse: 5’-TGGTTTCCTTCCTGGAGTTG-3’

HMGA2 forward: 5’-AGCGCCTCAGAAGAGAGGA-3’

HMGA2 reverse: 5’-AACTTGTTGTGGCCATTTCC-3’

### Protein expression by immunoblotting

Immunoblotting was performed using SDS-PAGE gels as previously described ^27^ using the following antibodies: anti-β-actin (Sigma, A5441, 1:10,000); anti-HRAS (Calbiochem, OP-23, 1:500); anti-NOTCH1 (Cell Signaling, 4380, 1:500); anti-HES1 (Cell signalling, 11988, 1:1,000); anti-FLAG (Cell Signaling, 2368, 1:1,000), anti-HMGA1 (Cold Spring Harbor Labs, #37, 1:1,000); anti-HMGA1 (Abcam, Ab4078, 1:1,000); anti-HMGA2 (Cold Spring Harbor Labs, #24, 1:1,000); anti-GFP (Clontech 632377, 1:1,000); anti-JAG1 (Cell Signaling, 2155, 1:1,000). Images of uncropped immunoblots are included in Supplementary Fig. 9.

### ATAC-seq

ATAC-seq samples were generated as previously described ^46^ using 100,000 IMR90 cells and 13-cycles of PCR amplification. Samples were size selected between 170 and 400bp (in order to isolate ‘nucleosome free’ and ‘mono-nucleosome’ fragments) using SPRIselect beads (B23319, Beckman Coulter) before single-end sequencing to generate 75-bp reads on the NextSeq-500 platform (Illumina).

### ChIP-seq

Chromatin immunoprecipitation (ChIP) was performed as previously described using 20 µg of sonicated chromatin ^76^ from Growing and RIS IMR90 ER:HRAS^G12V^ cells and 5µg of anti-H3K27Ac antibody (Clone CMA309 ^77^) and H3K4me1 antibody (Clone CMA302 ^77^). Libraries were prepared using the NEBNext Ultra II DNA Library Prep Kit for Illumina (37645, New England Biolabs) according to manufacturers instructions except that size selection was performed after PCR amplification using SPRIselect beads (B23319, Beckman Coulter). Samples were sequenced single-end using 50-bp reads on the HiSeq-2500 platform (Illumina).

### RNA-seq

RNA-seq data was generated from IMR90 ER:HRAS^G12V^ cells expressing a short-hairpin targeting the 3’ UTR of human *HMGA1* (RIS+shHMGA1). RNA was purified as above and quality checked using the Bioanalyser eukaryotic total RNA nano series II chip (Agilent). mRNA-seq libraries were prepared from 6 biological replicates of each condition using the TruSeq Stranded mRNA Library Prep Kit (Illumina) according to manufacturers instructions and sequenced using the HiSeq-2500 platform (Illumina).

### RNA-seq, ChIP-seq and ATAC-seq data analysis

#### RNA-seq analysis

Reads were mapped to the Human reference genome hg19 with the STAR (version 2.5.0b) aligner ^78^. Low quality reads (mapping quality less than 20) as well as known adapter contamination were filtered out using Cutadapt (version 1.10.0) (DOI:10.14806/ej.17.1.200). Read counting was performed using Bioconductor packages Rsubread and differential expression analysis with edgeR, ^79,80^. Differential expression analysis was carried out comparing the conditions against growing samples. Genes were identified as differentially expressed with a FDR cut-off of 0.01 and an absolute value of logFC (base 2 logarithm of the fold change) greater than 0.58.

#### ChIP-seq and ATAC-seq analysis

ChIP-seq and ATAC-seq reads were mapped to the human reference (hg19) with BWA (version 0.7.12) ^81^. A similar filtering was carried out using Cutadapt as described for RNA-seq data, and reads falling into the ‘blacklisted’ regions identified by ENCODE ^82^ were further removed. Average fragment size was determined using the ChIPQC Bioconductor package ^83^, and peak calling was performed with MACS2 (version 2.1.0) ^84^, using fragment size as an extension size (--extsize parameter). High confidence peak sets for each condition were identified separately, using only those peak regions that were present in at least two replicates.

#### Annotating peaks to genomic annotations

High confidence peak sets were mapped to genomic annotations or repeat regions using bedtools v2.26.0 ^85^. For the genomic annotations we used TSSs from the FANTOM database ^86^, repeats from repeatMasker (UCSC genome browser), and other genomic features (exons, introns, UTRs, etc.) were extracted from the UCSC Table Browser. The enhancers were identified based on our own H3K4me1 and H3K27ac histone mark ChIP-seq data sets; all regions that had peaks in both of these marks in either growing or RIS cells were considered as enhancers.

#### Intersecting consensus peaks and generating Venn diagrams

The Homer (v3.12) ^87^ command ‘mergePeaks’ with default settings and the output options ‘–venn’ and ‘–*prefix’* were used to generate values for plotting Venn diagrams and associated bed files for further analysis. Only literal overlaps (overlapping by 1bp) were considered. Venn diagrams were plotted using the R package ‘Venneuler’ (https://cran.rproject.org/web/packages/venneuler/index.html).

#### Calculating distances to Transcriptional Start Sites and GC percentages of peaks

To calculate the distance of consensus peaks from Transcriptional Start Sites (TSSs) and GC percentage of consensus peaks, the Homer (v3.12) ^87^ command ‘annotatePeaks.pl’ was used with default settings and the output option ‘*-*CpG’.

#### Gene ontology analysis

Consensus peaks within 300bp of a gene TSS were identified as described above. Gene Ontology (GO) analysis was performed using the GO Biological Process 2015 annotation provided on the web-tool ‘Enrichr’ ^88,89^ (http://amp.pharm.mssm.edu/Enrichr/).

#### Generation of normalised coverage files

A previously described trimmed-mean approach was used to generate scaling factors for each ATAC-seq condition relative to others ^47^. Briefly, we reasoned that the enrichment of reads within ATAC-seq peaks containing TSSs of genes that are both expressed (logCPM >mean logCPM) and have low variance between conditions (-0.14 < logFC < 0.14) by RNA-seq should not vary, unless there are differences in ATAC-seq sample quality, preparation or sequencing. By reanalysing our previously published IMR90 RNA-seq data ^15^ together with newly generated RNA-seq samples for RIS+shHMGA1 cells we identified 589 genes that fit these criteria. We counted the reads from the ATAC-seq samples that map to these specific genes using Rsubread, and computed scaling factors based on the mean counts for each condition separately. Normalised coverage files (bigWig) were generated by pooling reads from all of the replicates and applying the calculated scaling factors using the ‘genomecov’ function in bedtools, sorting the resulting normalised bedGraph files and then converting them to bigWigs using the ‘bedGraphToBigWig’ function from UCSC.

#### Generation of clustered heatmaps

Heatmaps were generated using normalised coverage of peaks (+/-1kb) representing novel NFRs (consensus peaks detected in any sample excluding peaks detected in the growing IMR90 sample) with *k-means* clustering using the deepTools package ^90^.

#### TCGA analysis

We analysed expression levels of NOTCH-associated genes in the publically available RNA sequencing data generated by the TCGA Research Network: http://cancergenome.nih.gov/ ^55^. Computational analysis and statistical testing of the NGS data was conducted using the R statistical programming language ^91^. Filtered and log 2 normalised RNA expression data along with all available clinical data were downloaded from the GDAC firehose database (run: stddata 2015_06_01) for each gene of interest from the relevant cancer-specific collections.

Correlation testing for associations between expressed genes was performed using the cor.test function in R to calculate the Pearson's product moment correlation coefficient and test for significant deviation from no correlation. Plotting of TCGA data was performed using the ggplot2 R package ^92^. Survival analysis was performed using the survminer and survival ^93^ R packages. Kaplan-Meier estimated survival curves were constructed using the TCGA clinical data. Statistical testing of differences between survival curves used the G-rho family of tests, as implemented in the survdiff function of the survival package.

## ACKNOWLEDGEMENTS

We thank all members of the Narita laboratory for helpful discussions, M. de la Roche for reagents, and staff at the Genomics, Light Microscopy, Biorepository, Flow Cytometry and Research Instrumentation core facilities at the Cancer Research UK Cambridge Institute for technical support. The University of Cambridge, Cancer Research UK and Hutchison Whampoa supported this work. The M.N. and S.B. laboratories are funded by a Cancer Research UK Cambridge Institute Core Grant (C14303/A17197). M.H. is supported by a CRUK Clinician Scientist Fellowship (C52489/A19924). R.H.-H. is funded by an EMBO Long-Term fellowship. S.A.S. and D.B. were supported by Medical Research Council core funding.

## AUTHOR CONTRIBUTIONS

A.J.P., M.H. and R.H.-H. designed experiments, performed experiments and analysed data. D.B., S.A.S. and A.J.P. analysed sequencing data. S.S. analysed TCGA data. H.K. provided the antibody for H3K27Ac and H3K4me1 ChIP-seq. S.B., S.A.S., H.K. and M.N. supervised experiments and interpreted the data. M.N., M.H. and A.J.P. conceived the project. M.N. and A.J.P. wrote the manuscript with contribution and review from all authors.

## COMPETING FINANCIAL INTERESTS

The authors declare no competing financial interests.

---

**Supplemental Figure 1.**
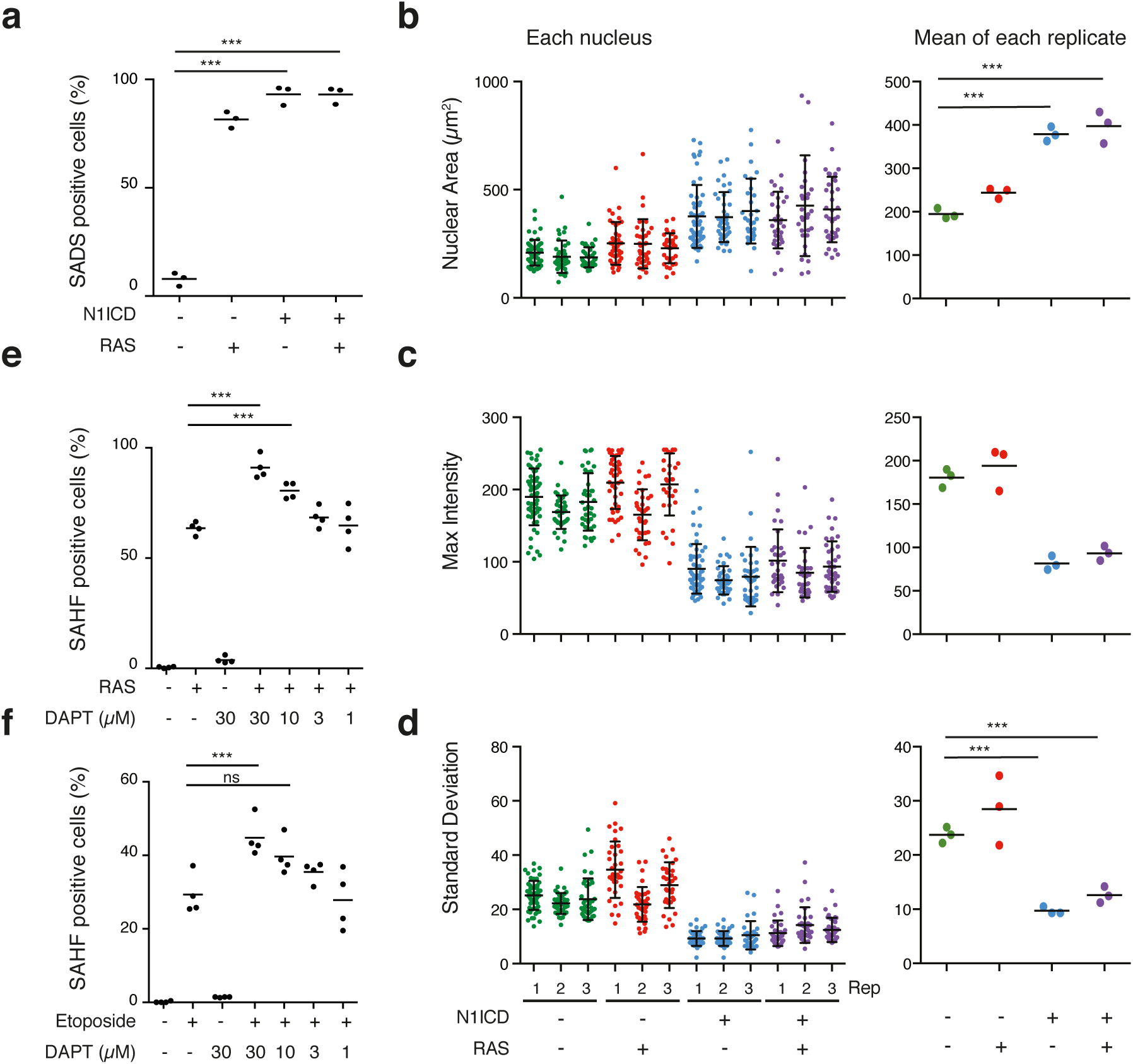
NOTCH1 signalling has a chromatin ‘smoothening’ effect that strongly blocks SAHFs **(a-d)** IMR90 ER:HRAS^G12V^ cells infected with control vector or FLAG-N1ICD ± 4OHT. **(a)** Percentage of SADS positive cells. **(b-d)** Quantification of nuclear area **(b)**, maximum pixel intensity **(c)** and standard deviation **(d)** of signal from at least 30 DAPI stained nuclei per biological replicate for the conditions indicated. Data for individual nuclei (left, mean ± s.d is plotted) and means of each replicate (right) are presented. n=3 biologically independent replicates each. **(e, f)** Quantification of SAHF positive cells in IMR90 ER:HRAS^G12V^ cells ± 4OHT **(e)** or IMR90 cells treated with etoposide **(f)** and different concentrations of DAPT. n = 4 biologically independent replicates each; Statistical significance calculated using one-way ANOVA with Tukey’s correction for multiple comparisons.

**Supplemental Figure 2.**
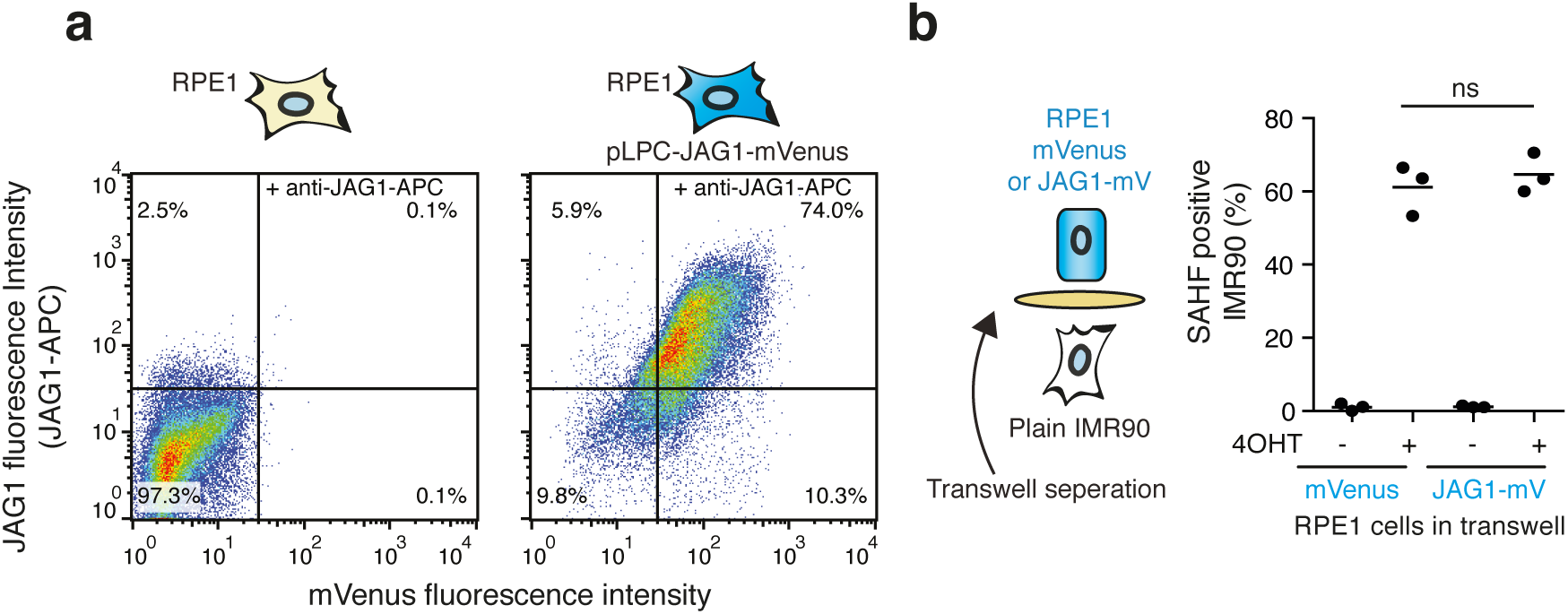
Ectopic JAG1 expressing RPE1 cells require cell-cell contact to repress SAHFs in RIS cells **(a)** RPE1 JAGGED1-mVenus cells analysed for cell surface expression of JAG1 by flow cytometry (n=1). **(b)** Quantification of SAHF positive IMR90 ER:HRAS^G12V^ cells cultured in a transwell dish with the indicated RPE1 cells for 6 days +4OHT (n=3)

**Supplemental Figure 3.**
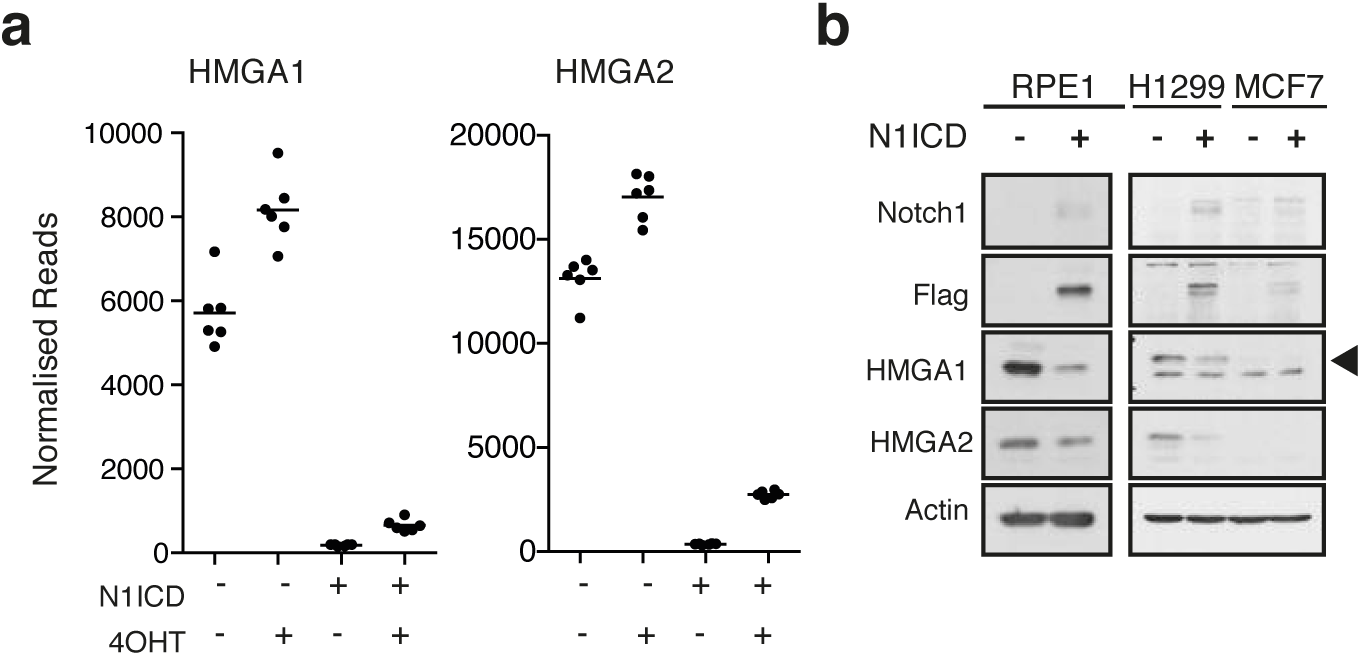
NOTCH signalling represses HMGAs **(a)** Re-analysis of previously published RNA-seq data ^15^ for the cell conditions and genes indicated. **(b)** Immunoblotting of RPE1, H1299 and MCF7 cells expressing FLAG-N1ICD for the proteins indicated.

**Supplemental Figure 4.**
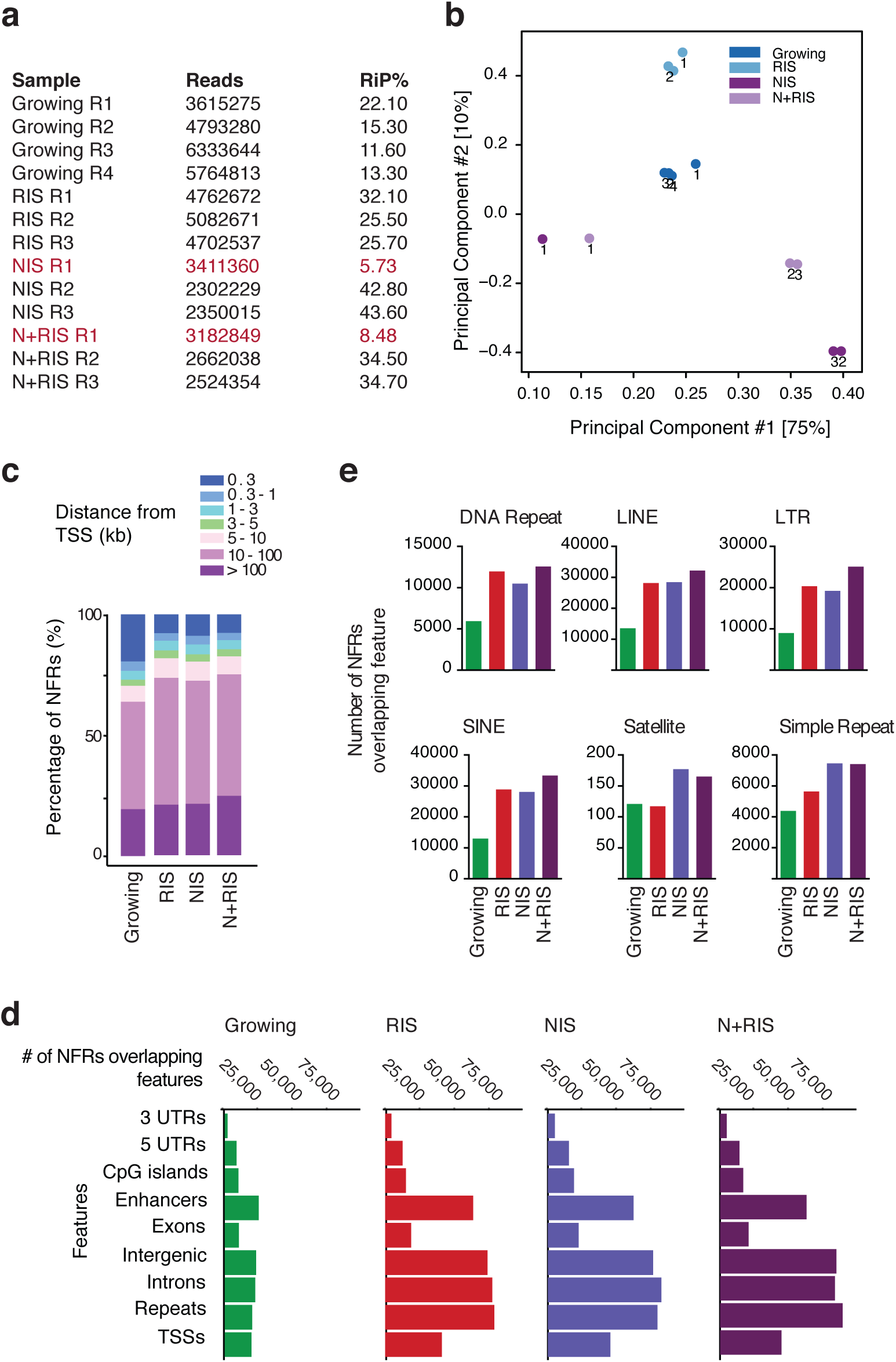
ATAC-seq data is of high quality and aligns to gene distal elements **(a)** Number of valid reads and percentage of reads in peaks (RiP%) for the ATAC-seq samples indicated. **(b)** Unbiased PCA analysis of raw ATAC-seq samples. **(c)** Percentage of high-confidence NFRs (present in at least two replicates) within the indicated distances of a transcriptional start site (TSS). **(d, e)** High-confidence NFRs in each cell condition annotated to genomic annotations **(d)** and repeat types **(e)**.

**Supplemental Figure 5.**
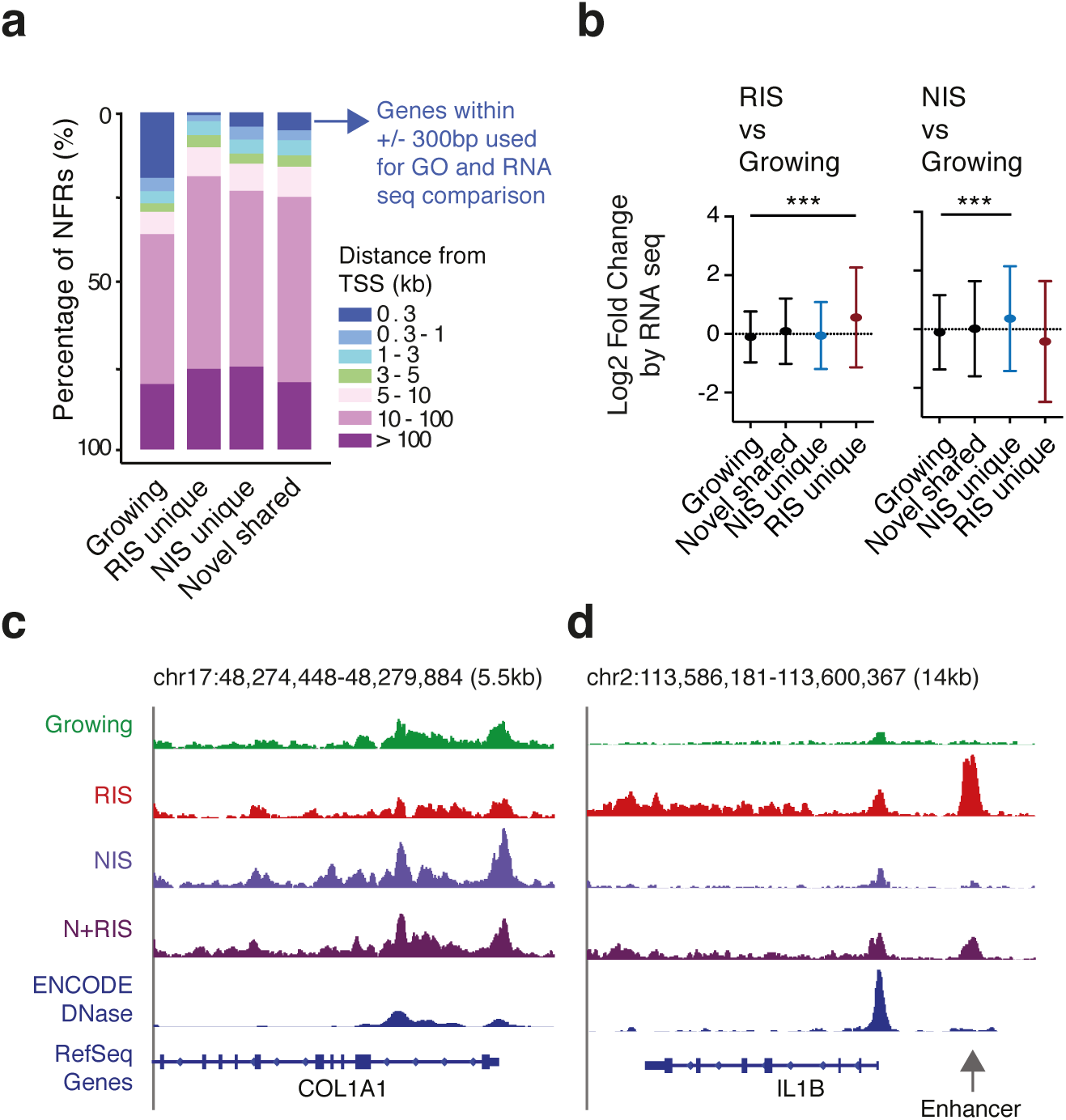
NFRs reflect gene transcription in RIS and NIS cells **(a)** Percentage of NFRs from the indicated subsets (defined in Fig. 4e) within the indicated distances of a transcriptional start site (TSS). **(b)** NFRs within the indicated subsets (defined in Fig. 4e) were annotated to genes if within 300bp of a TSS. The average log2 fold expression change of genes in each subset in RIS and NIS cells in comparison to growing cells is plotted. Previously published RNA-seq data was used ^15^. Values are mean ± s.d. Statistical significance calculated using one-way ANOVA with Tukey’s correction for multiple comparisons; *P ≤ 0.05, **P ≤ 0.01, ***P ≤ 0.001. **(c, d)** Representative genome browser images of normalised coverage data for the conditions indicated. The COL1A1 locus **(c)** is transcriptionally activated in NIS and RIS+N1ICD. The IL1B gene **(d)** is transcribed in RIS cells only, potentially from an upstream enhancer (indicated)

**Supplemental Figure 6.**
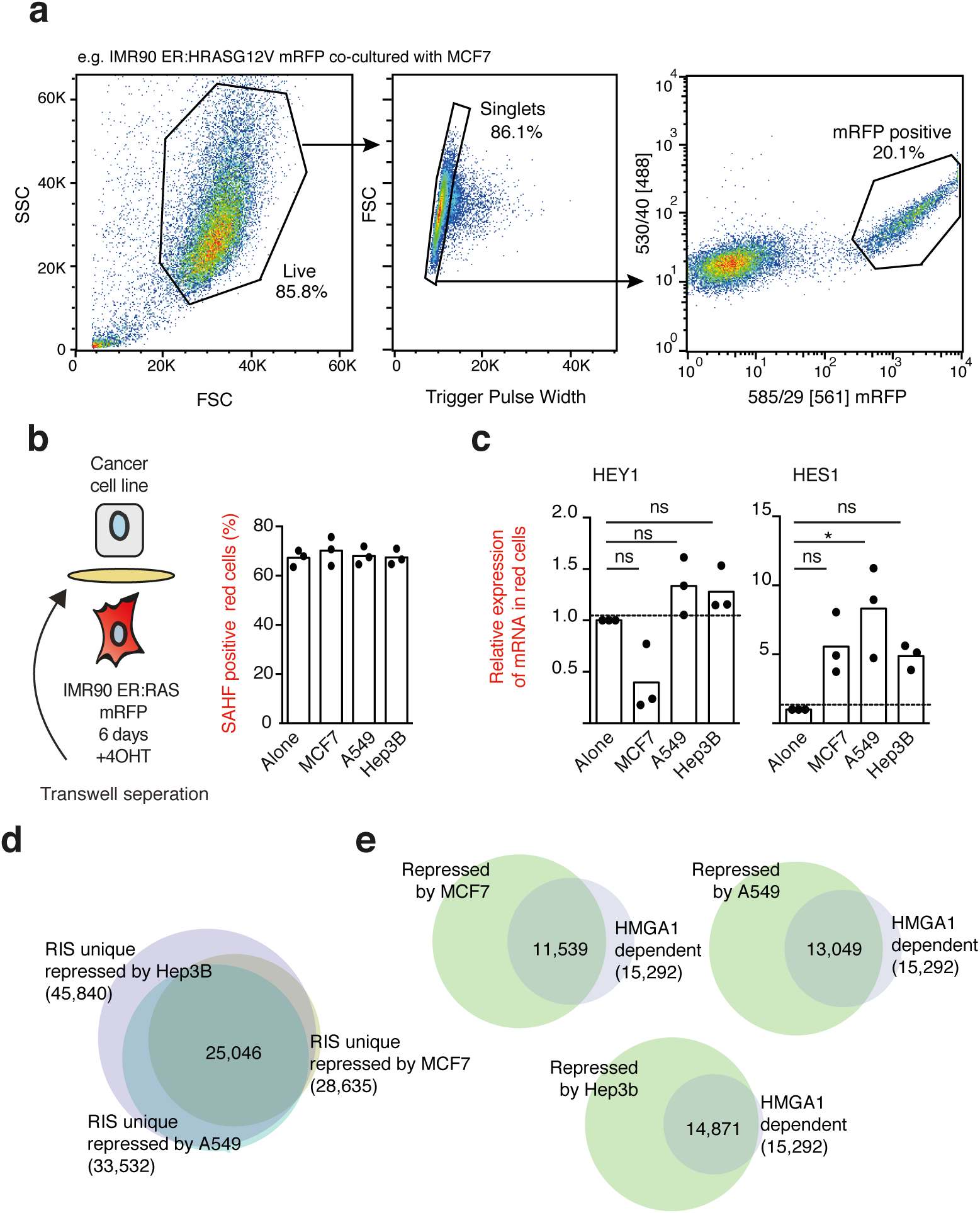
Tumour cells repress RIS specific NFRs in adjacent fibroblasts in a cell-contact dependent manner. **(a)** Example of flow-sorting strategy for the isolation of mRFP positive cells from co-cultures between tumour cell lines and IMR90 ER:HRASG12V cells expressing mRFP1. **(b)** Quantification of SAHF positive IMR90 ER:HRAS^G12V^ cells cultured in a transwell dish with the indicated tumour cell lines cells for 6 days +4OHT. **(c)** qRT-PCR of mRNA isolated from flow sorted IMR90 ER:HRAS^G12V^ mRFP1 cells cultured with tumour cell lines +4OHT for 6 days relative to cells cultured alone (as described in Fig. 6c). **(b, c**) n = 3 biologically independent replicates. Statistical significance calculated using one-way ANOVA with Tukey’s correction for multiple comparisons; *P ≤ 0.05, **P ≤ 0.01, ***P ≤ 0.001. **(d, e)** Venn-diagrams showing literal overlap between NFRs identified in the subsets indicated. Overlaps are between ‘RIS specific NFRs’ (defined in Fig 3e) that are not detected in the RIS+MCF7, RIS+A549 and RIS+Hep3B ATAC-seq samples **(d)** or between these and ‘HMGA1 dependent NFRs’ defined in Fig. 5a **(e)**.

**Supplemental Figure 7.**
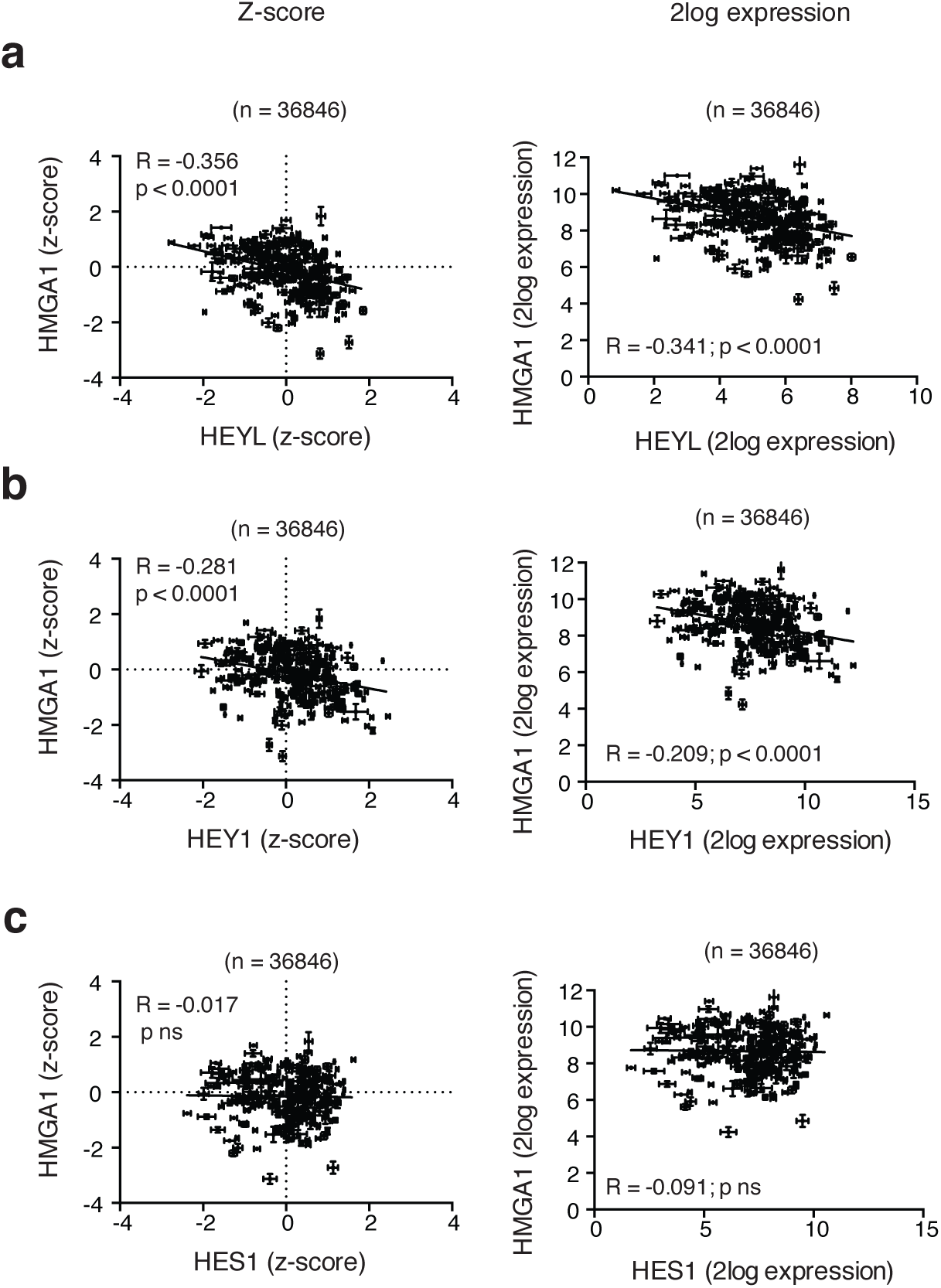
HMGA1 gene expression anti-correlates with canonical NOTCH target genes **(a-c)** The R2 database (http://r2.amc.nl) was used to compare the microarray determined mRNA expression of HMGA1 with HEYL **(a)**, HEY1 **(b)** and HES1 **(c)**.

**Supplemental Figure 8.**
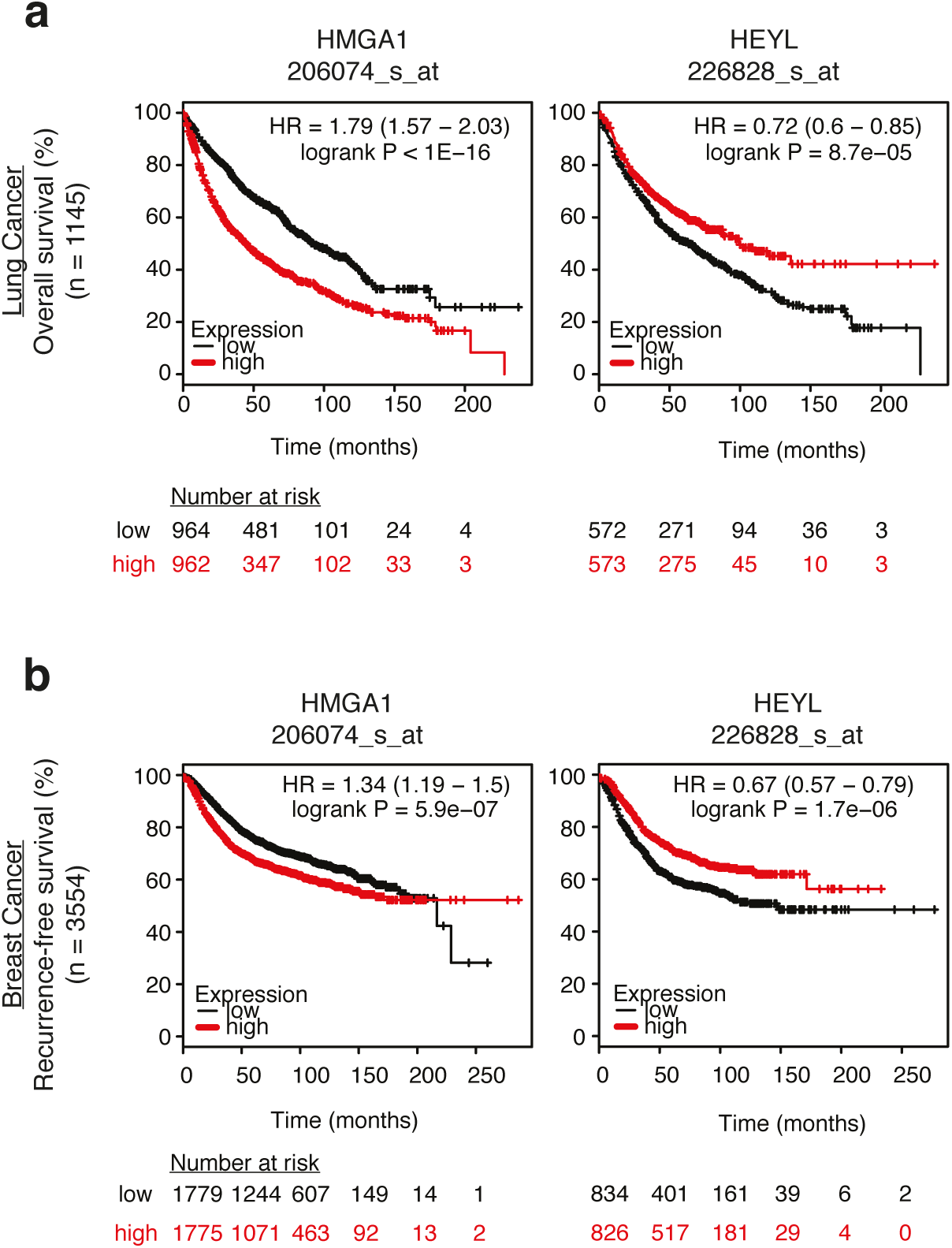
HEY-L and HMGA1 gene expression is prognostic in lung and breast cancers. **(a, b)** Kaplan-Meier plots of overall survival of 1145 patients with lung cancer **(a)** and recurrence-free survival of 3554 patients with breast cancer **(b)** stratified based on microarray determined mRNA expression of HMGA1 (left panels) and HEYL (right panels).

